# The desert green algae *Chlorella ohadii* thrives at excessively high light intensities by exceptionally enhancing the mechanisms that protect photosynthesis from photoinhibition

**DOI:** 10.1101/2021.02.08.430232

**Authors:** Guy Levin, Sharon Kulikovsky, Varda Liveanu, Benjamin Eichenbaum, Ayala Meir, Tal Isaacson, Yaakov Tadmor, Noam Adir, Gadi Schuster

## Abstract

Although light is the driving force of photosynthesis, excessive light can be harmful. One of the main processes that limits photosynthesis is photoinhibition, the process of light-induced photodamage. When the absorbed light exceeds the amount that is dissipated by photosynthetic electron flow and other processes, damaging radicals are formed that mostly inactivate photosystem II (PSII). Damaged PSII must be replaced by a newly repaired complex in order to preserve full photosynthetic activity. *Chlorella ohadii* is a green micro-alga, isolated from biological desert soil crusts, that thrives under extreme high light and is highly resistant to photoinhibition. Therefore, *C. ohadii* is an ideal model for studying the molecular mechanisms underlying protection against photoinhibition. Comparison of the thylakoids of *C. ohadii* cells that were grown under low light versus extreme high light intensities, found that the alga employs all three known photoinhibition protection mechanisms: *i)* massive reduction of the PSII antenna size; *ii)* accumulation of protective carotenoids; and *iii)* very rapid repair of photo-damaged reaction center proteins. This work elucidated the molecular mechanisms of photoinhibition resistance in one of the most light-tolerant photosynthetic organisms and shows how photoinhibition protection mechanisms evolved to marginal conditions, enabling photosynthesis-dependent life in severe habitats.

**One Sentence Highlight:** Analysis of the photosynthetic properties of a desert algae that thrives at extreme high light intensities revealed protection from photoinhibition driven by the remarkable enhancement of three protection mechanisms.

## Introduction

Oxygenic photosynthesis, which takes place in cyanobacteria, algae and plants, provides all of the food and is the source of carbon-based fuels on earth (Blankenship, 2014; Barber, 2009). In eukaryotes, two multi-subunit pigment-protein complexes, photosystem II and I (PSII and PSI, respectively), that are located in the thylakoid membranes of chloroplasts, absorb light, converting this excitation energy into chemical energy stored in chemical bonds. While light is the driving force of photosynthesis, excessive light can be harmful, with the threshold of the damaging light intensity depending on the organism’s natural environment. Photoinhibition (PI) occurs when the absorbed light energy exceeds the capacity of the plant to utilize it via electron transfer, photochemical quenching or non-photochemical quenching pathways (Gururani *et al*., 2015). PSII is the most sensitive complex in the thylakoid to photodamage, while PSI is relatively stable (Nishiyama and Murata, 2014; Li *et al*., 2009; Lu, 2016; Adir *et al*., 2003; Tikkanen *et al*., 2014). Excessive light promotes formation of reactive oxygen species (ROS), which, in turn, react with PSII components. Once PSII proteins are damaged, the complex becomes inactive and a repair cycle is initiated, during which replacement of damaged proteins occurs and PSII activity is restored (Nishiyama and Murata, 2014; Li *et al*., 2009; Lu, 2016; Keren and Krieger-Liszkay, 2011; Li *et al*., 2018; Cruz *et al*., 2005; Erickson *et al*., 2015; Chotewutmontri and Barkan, 2020). PI occurs when the rate of damage caused by oxidative stress exceeds the rate of PSII repair (Vecchi *et al*., 2020; Liu *et al*., 2019; Rochaix, 2020). In many photosynthetic organisms, there are special stress-related light-harvesting II (LHCII) subunits, termed LHCSR, that accumulate high-light conditions and direct the absorbed light to non-photochemical processes that reduce the production of oxygen radicals (Erickson *et al*., 2015; Larkum *et al*., 2020; Rochaix and Bassi, 2019; Girolomoni *et al*., 2019; Pinnola, 2019). Nevertheless, the high-light tolerance green algae *Chlorella ohadii*, which is the research topic of this study, lacks the LHCSR proteins.

The second mode of defense against PI in photosynthetic organisms involves a decrease in PSII antenna size. This is achieved by phosphorylation-activated disconnection of LHCII from PSII, followed by a decline in chlorophyll *b* (chl *b*) content and a reduction of LHCII expression, thereby curbing the uptake of excess light energy (Rochaix and Bassi, 2019; Ananyev *et al*., 2017; Vecchi *et al*., 2020; Nicol *et al*., 2019; Croce, 2020). Overall, the change leads to more efficient photosynthetic activity due to reduced generation of ROS, and maintains a high steady state level of photosynthesis. The latter occurs since the PSII repair cycle is continuously replacing the damaged reaction center (RC) proteins (Rochaix, 2020; Kolodny *et al*., 2021; Wobbe *et al*., 2016; Friedland *et al*., 2019; Stefano *et al*., 2014).

The third major mechanism for protection from PI involves the accumulation of carotenoids in the vicinity of the chlorophylls of PSII and LHCII. These carotenoids, such as lutein and β-carotene, quench the energy of the excited chlorophylls, preventing the formation of harmful radicals (Bassi and Caffarri, 2000; Murchie and Ruban, 2019; Leonelli *et al*., 2017). In the underlying “xanthophylls cycle”, violaxanthin is converted in high light (HL) to antheraxanthin, which is further converted to zeaxanthin. Zeaxanthin accumulation in the PSII quenches the photo-excited chlorophyll radicals to produce heat (Niyogi, 1999; Nicol *et al*., 2019; Leonelli *et al*., 2017; Vecchi *et al*., 2020; Nishiyama and Murata, 2014).

*Chlorella ohadii* is a green micro-alga isolated from biological soil crusts in the Negev desert of Israel (Treves *et al*., 2016; Treves *et al*., 2013). Characterized by subfreezing temperatures in winter and extremely high temperatures in summer, the desert is a very harsh habitat. The dry and arid environment is accompanied by intense sunlight during the long desert summer, forcing photosynthetic organisms to develop exceptional mechanisms to avoid PI. Previous studies have suggested some of the adaptations that enable *C. ohadii* to thrive under these extreme conditions (Treves *et al*., 2016). These include a short generation time (Treves *et al*., 2017), significant PSII-cyclic electron flow under HL intensities (Ananyev *et al*., 2017), maintained maximum PSII efficiency in HL intensities of 3500 µE m^-2^s^-1^ (Treves *et al*., 2016; Treves *et al*., 2020), as well as multiphasic growth controlled by metabolic shifts (Treves *et al*., 2017). Considering its remarkable PI resistance and record growth rate, *C. ohadii* can serve as an ideal model organism for the study of PI resistance (Vecchi *et al*., 2020; Leister, 2019).

In this work, we compared the thylakoids structure and photosynthetic activities of *C. ohadii* grown under extreme HL versus low light (LL) intensities. When grown under an extreme HL intensity, an exceptionally marked enhancement of the three major mechanisms of protection from PI to marginal states of use was noted. More specifically, the peripheral PSII antenna was nearly entirely eliminated, the xanthophyll cycle was highly activated and the PSII repair cycle replaced photo-damaged proteins faster than any other known photosynthetic eukaryote. Therefore, the extreme resistance to PI was evolutionarily achieved in this organism by recruiting all three known major anti-PI mechanisms at maximal efficiency.

## Results and Discussion

### Differences in the photosynthetic parameters between HL and LL cells

In order to study changes in thylakoids of *C. ohadii* cells that thrive under extreme HL as compared to those that grow under LL intensities, we compared the photosynthetic properties and thylakoids of cells growing at these conditions. *C. ohadii* cells were grown for 24 h at low light of 50 μE m^−2^ s^−1^ (LL), or high light intensity of 2500 μE m^−2^ s^−1^ (HL), the latter being approximately four times the light intensity of a sunny summer noon in their living habitat and the intensity of saturating photosynthesis. In order to prevent self-shading of the cells, the OD_750_ was maintained below 0.7 for the entire growth period (about 6 generations) by diluting the culture. *C. ohadii* is one of the most rapidly reproducing photosynthetic eukaryotes, with a reported division time as rapid as 2 h, but which was found to be about 4 h in our growth system (Table 1) (Treves *et al*., 2017; Treves *et al*., 2020). Under both LL and HL conditions, the cells grew rapidly, displaying the same generation time of about 4 h, demonstrating the high tendency of *C. ohadii* to thrive under extreme HL (Table 1 and Figure S1). Similar results were obtained with *Chlorella sorokiniana*, a species closely related to *C. ohadii* (not shown).

**Table 1.**
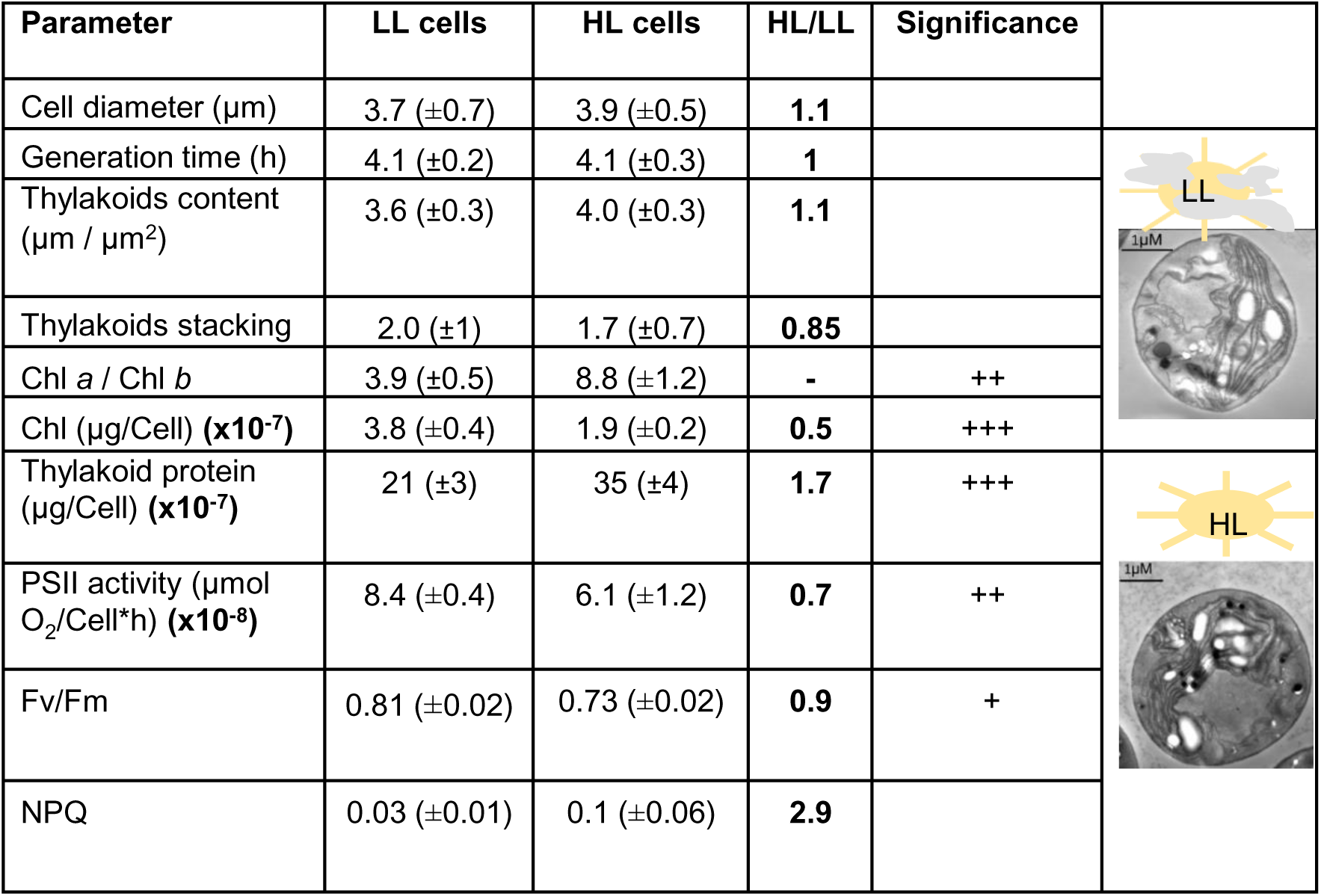
Differences between HL- and LL-grown cells of *C. ohadii*. Cell size, thylakoid content, degree of stacking, chlorophyll content, thylakoid protein content and PSII activity were determined and calculated per *C. ohadii* cell of HL and LL grown cells. Cell size was determined by measuring the diameter of cells of similar sections in electron microscope (EM) images (Fig S2). The amount of thylakoid content per cell was determined by measuring the length and stacking degree in EM images (see Figs S2 and S3). PSII activity was determined as the rate of O_2_ evolution using DCBQ as an electron acceptor and from the fluorescence kinetics of PSII. NPQ was analyzed from the fluorescence kinetics data of PSII. Significance of at least three independent experiments was determined using two-tailed distribution students t test. **+** = p<0.05, **++**=p<0.005, **+++**=p<0.0005. Chl-Chlorophyll. For full details, see Table S1.

HL and LL thylakoids differed in their protein and chlorophyll content, which are typically used for normalization when comparing thylakoids from different cell types. Therefore, we searched for a parameter that was comparable and could be used for normalization of the photosynthetic measurements. We found that the cell size and number of thylakoids were very similar under both growth conditions (Table 1). Close examination of numerous electron microscope images identified very similar length and stacking degree of thylakoids in LL and HL cells (Table 1, Figures S2 and S3), in contrast to previous reports which showed that the stacking degree is significantly increased in *C. ohadii* cells exposed to a similar HL intensity (Treves *et al*., 2016; Treves *et al*., 2020). Therefore, since the cell size and the length and number of thylakoids were similar in HL and LL cells, all the measured parameters were normalized to the corresponding amount in identical cell numbers.

Chlorophyll (chl) *b* content was significantly reduced in HL thylakoids, leading to an increase in the chl *a/b* ratio from 4.0-5.0 in LL to 8.8 in HL (Table 1). Chl *b* is mostly present at the PSII light harvesting antenna (LHCII), and, to a lesser extent, at the PSI antenna (LHCI). Its depletion under HL conditions suggested a significant reduction in LHCII. Concomitantly, the total chlorophyll content per HL cell was reduced to about half of that measured in the LL cells, while the total protein content increased 1.7-fold in HL thylakoids (Tables 1 and S1 and Figure S4). While both HL and LL thylakoids displayed high oxygen evolution activity of PSII, it was about 30% lower in the HL cells as compared to the LL cells (Tables 1 and S1). The Fv/Fm variable fluorescence, another measure of PSII activity, was only 10% lower in HL cells as compared to LL cells (Table 1 and Figure 1C). These measurements indicate that HL cells were highly active in linear electron flow photosynthesis and were not severely photoinhibited, clearly showing the ability of HL cells to thrive and rapidly reproduce under such extreme HL intensities. Although the non-photochemical chlorophyll fluorescence quenching (NPQ) was very low as compared to plants and other green algae, presumably due to the lack of the stress-related LHC (LHCSR) proteins in this strain (Treves *et al*., 2016), it was three-fold higher in HL as compared to LL cells (Table 1 and Figure 1). The HL cells and their thylakoids were yellowish and olive-colored following HL growth, and deep green under LL, reflecting the significant accumulation of carotenoids in the former (see below).

**Figure 1.**
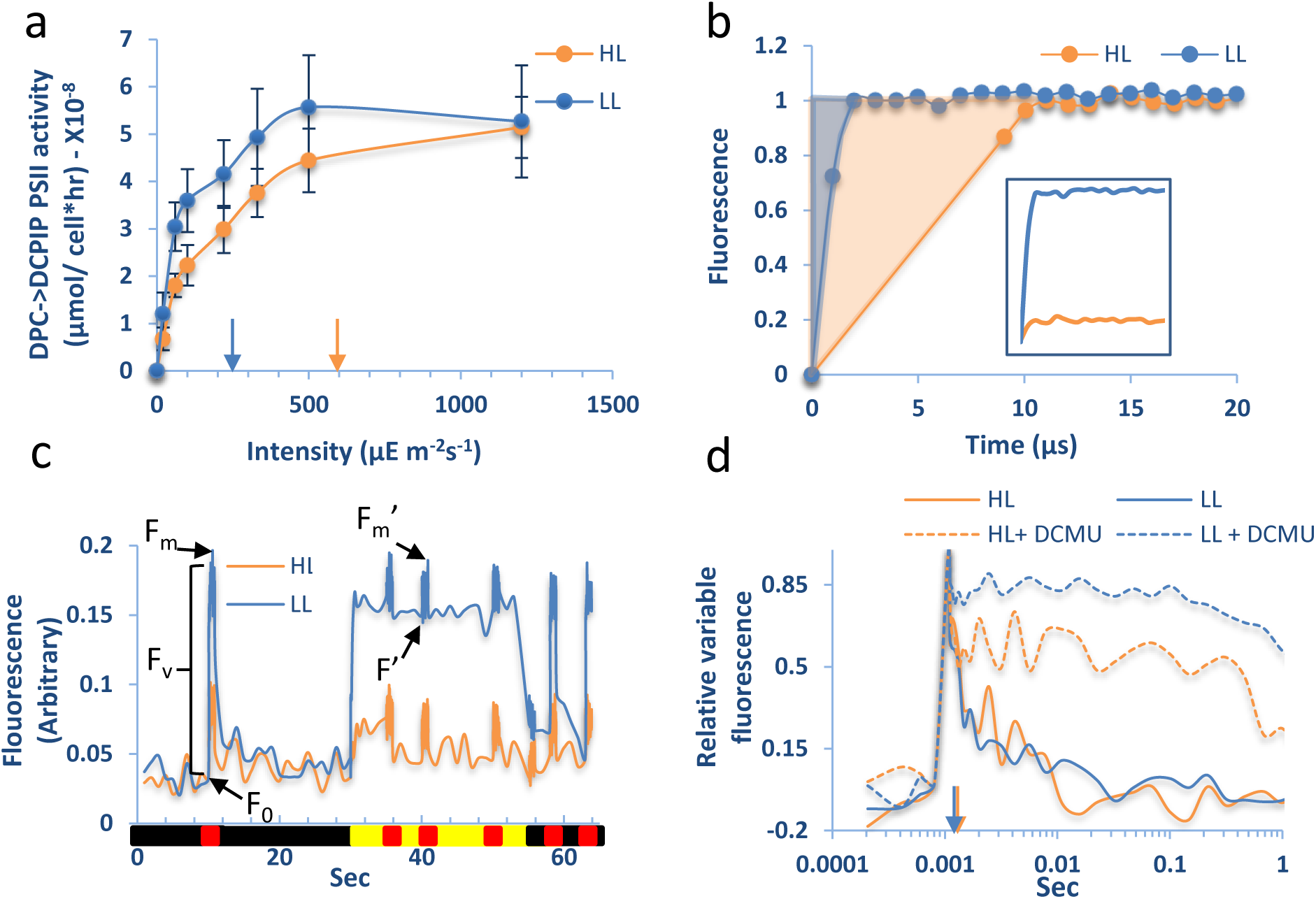
Photosynthetic activities and fluorescence of HL and LL cells reveal reduced PSII antenna size and similar linear electron flow. **a.** Saturation curves of the light intensity-associated PSII activities in LL and HL thylakoids. Arrows indicate the light intensities in which half saturation of PSII activity were detected. Error bars present the standard deviation (SD) of at least three experiments. **b.** Functional PSII antenna size decreases in HL. Typical fluorescence induction traces of LL (blue) and HL (orange) in limited light and in the presence of DCMU. The curves were normalized to F_m_ and F_0_, as described in the Methods. The area above the trace (indicated in light-blue and light-orange colors) is reciprocally proportional to the functional PSII antenna size, according to the slopes of several measurements, as shown in Figure S5b. The same curve without normalization is shown in the inset of the panel, and shows the strong quenching of F_m_ in HL. **c**. Kinetics of the fluorescence induction traces of PSII of dark-adapted LL and HL cells normalized to F_0_. The fluorescence signals were measured in an equal number of cells. The illumination time frame throughout the measurement is indicated on the X axis: black: dark, red: high-intensity saturating pulse, yellow: actinic light. **d.** Fluorescence traces of the Q_A_ relaxation kinetics measured the Q_A_ forward electron flow rate in the absence (full lines), and compared it to the Q_A_ back electrons flow rate with the addition of DCMU (dashed lines). Although the intensity of the PSII fluorescence was significantly lower in HL cells, F_0_ and F_m_ were normalized to zero and one, respectively, in order to detect the time of relaxation of the signal to 50% (indicated by an arrowhead). The rate of forward electron flow was similar to that measured in LL cells, as shown by the similar Fv/Fm and the half time of the Q_A_ relaxation time. In panels b-d, representative results of a single experiment are presented, of which at least three were conducted and showed the same results.

### HL thylakoids lose nearly the entire trimeric LHCII antenna

The dramatic reduction in the amount of chl *b* in HL thylakoids may be due to depletion of this pigment from the PSII antenna or the lack of the LHCII. In order to measure the size of the PSII functional antenna, the saturation curve of PSII activity was analyzed under increasing light intensities. A significant increase in the slope was observed at low light intensities of up to 100 μE m^-2^s^-1^ in LL as compared to HL thylakoids (Figure 1a). The level of PSII functioning in the LL thylakoids reached saturation at 500 μE m^-2^ s^-1^, while in HL thylakoids saturation was not reached, even at the maximal intensity tested. The differences between the slopes of the curves at the lowest light intensities indicated an approximate 40% decrease in the functional size of LHCII in HL, as compared to LL cells. A second method is to analyze the area above the fluorescence rise kinetic traces taken under low light intensities and in the presence of the electron transfer inhibitor, 3-(3,4-dichlorophenyl)-1,1-dimethylurea (DCMU). Following normalization of F_m_ and F_0_ and correction for F_0_ parameters, as described by Tian *et al*. (Tian *et al*., 2019), the relative PSII antenna size was determined (Figure 1b and S5). The results revealed 80% decrease in the functional size of LHCII in HL, as compared to LL cells. Even though the modes of reduction of the functional LHCII as determined by the two systems were not identical, probably due to the remarkable differences in the methods, both indicated a significant and considerable reduction in functional LHCII content. The reduction in LHCII size was further corroborated by low temperature (77K) fluorescence measurements of LL and HL cells. When normalized to the fluorescence of the LHCI at 723 nm, assumed to be invariant, the relative intensities of the fluorescence of the LHCII and PSII-RC in HL cells at 686 nm and 696 nm were significantly lower than in LL cells (Figure S5). Notably, the PSI peak at 708 nm was significantly increased in HL cells. The nature of this increase is still unknown. Quenching of LHCII-PSII chlorophyll fluorescence could be the result of a reduction in the antenna size, as shown above for PSII, and of the accumulation of carotenoids that perform NPQ.

Further evidence for the simultaneous reduction in fluorescence properties of the PSII antenna chlorophylls in HL cells and conservation of high linear electron flow activity, was obtained by measuring both the kinetics of fluorescence induction and the Q_A_ relaxation parameters (Figures 1c and 1d). In both measurements, the relative fluorescence of the PSII chlorophylls was much lower in HL cells, indicating the presence of an efficient quenching mechanism. It should be noted that the graph of the Q_A_ relaxation kinetics of the HL cells was expanded and normalized to that of LL in order to determine the half-life of the relaxation time of the forward electron flow (Figure 1d). Since *C. ohadii* lacks the LHCSR complexes, the quenching can be ascribed to the significant accumulation of carotenoids, and/or the activity of psbS, which is overexpressed in high light (Girolomoni *et al*., 2019; Pinnola, 2019; Larkum *et al*., 2020; Rochaix and Bassi, 2019; Nicol *et al*., 2019; Głowacka *et al*., 2018; Tian *et al*., 2019; Treves *et al*., 2020). Even though the fluorescence signal was quenched in HL cells, the rate of linear PSII electron flow remained high, as manifested by the similar Fv/Fm, Q_A_ relaxation half-life time and PSII electron transport rates, indicating the resistance of *C. ohadii* to PI while growing for many generations at HL.

In order to study possible changes in the structures of PSII and PSI upon exposure to LL versus HL, the photosynthetic complexes were extracted using mild detergent treatments, and fractionated by sucrose density gradient or native PAGE, which was further analyzed by denaturing SDS-PAGE. The results of these analyses, normalized either to equal amounts of chlorophyll or to the amount of the different complexes per HL or LL cells, disclosed that HL thylakoids displayed a marked reduction in PSII-LHCII complexes and the trimeric LHCII (Figures 2a, 2b). Analyzing the profile of the complexes separated by the sucrose density gradient clearly disclosed the extensive reduction in the amount of the PSII-LHCII and the corresponding accumulation of the PSII core in HL (Figure 1a). Note that thylakoid extracts of equal amounts of chlorophyll were fractionated in this assay in order to better observe the complexes of HL thylakoids. This resulted in twice the number of thylakoids of HL as compared to the quantities seen for samples normalized by complexes per cell. When comparing the gradients from these cells, the position of the PSI-LHCI complex remained constant, while the LHCII of HL was more disperse and was found at a lower sucrose density (Figure 1a).

**Figure 2.**
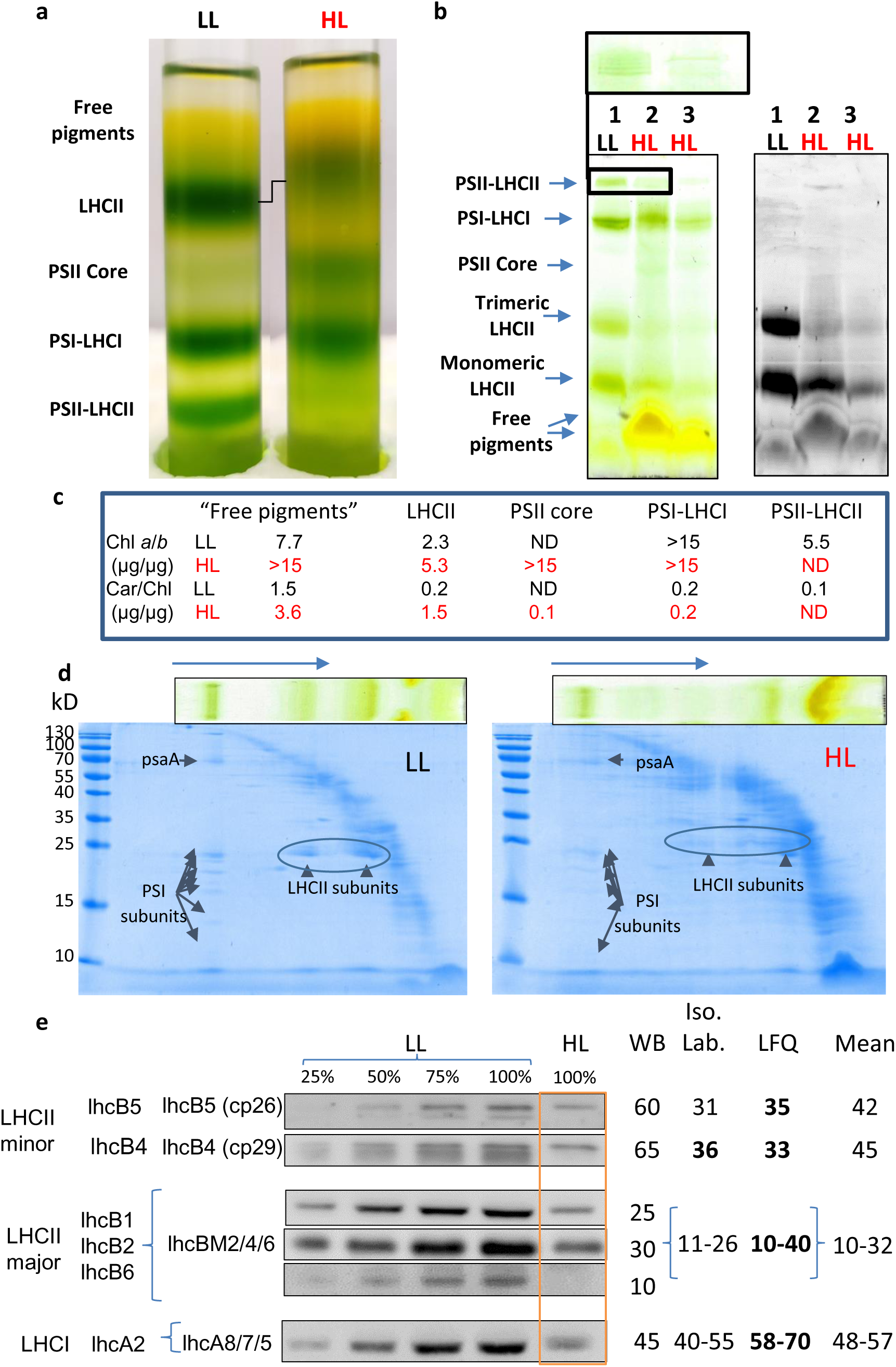
LHCII is downregulated in HL cells. **(a)** Photosynthetic complexes were extracted from thylakoids of HL and LL cells and fractionated by sucrose density gradients. Thylakoid extracts containing equal amounts of chlorophyll were loaded; therefore, each complex observed in HL represents the quantity from twice the amount of cells/thylakoids as compared to LL. **(b)** Analysis of the chlorophyll-protein complexes by non-denaturing native PAGE. Thylakoids containing 8 μg chlorophyll from LL cells (1) and 8 μg (2) or 4 μg (3) chlorophyll for HL cells were loaded in order to observe the amount of complexes at equal chlorophyll levels (1 and 2) and to normalize the relative amounts of LL and HL complexes per cell (1 and 3). A magnification of the PSII-LHCII region, indicated by a black box, is presented on the top to visualize the different complexes of this kind. Note the considerable accumulation of non-complexed pigment in the gel-front of the lane of the HL sample. Fluorescence of the same native PAGE is shown to the right. The LHCII but not the PSII or PSI-LHCI complexes display fluorescence emission. **(c)** The chlorophyll (Chl) *a/b* and total carotenoids (Car) ratios (µg/µg) in the photosynthetic complexes and in the “free pigments” measured in the sucrose density gradient separation of LL and HL thylakoids shown in panel a. ND-not detected. **(d)** Second-dimension gel analysis of the native PAGE. The non-denaturing gels of LL (left panel) and HL (right panel) thylakoids were denatured and further fractionated by denaturing SDS-PAGE. The PSI and LHCII proteins are indicated. (**e**) Immunoblotting was performed to assess the downregulation of specific LHC subunits. To identify the corresponding *C. ohadii* LHC subunit, antibodies produced against Arabidopsis LHC proteins were used; see experimental section and Table S3. The legends from left to right indicate the photosynthetic complex, the name of the Arabidopsis protein to which the antibody was raised and the possible *C. ohadii* homologs. It was impossible to definitively define a single *C. ohadii* homolog of the Arabidopsis LHC proteins and, therefore, the three candidate proteins are indicated (see Table S3). (**e, right panel**). Values in the table indicate the relative abundance of the subunit in HL compared to LL thylakoids, as measured via western blot, isotope labelling mass spectrometry and label-free quantification (LFQ) mass spectrometry (Table S2). The mean amount protein measured by the three methods is indicated in the right lane. The two proteomic methods were normalized to demonstrate changes in the levels of the specific protein per cell. This was done by dividing the LL amounts by 1.7, since there are x1.7 fewer proteins in LL as compared to HL thylakoids (Table 1). Bold numbers indicate statistically significant changes (For LFQ: FDR<0.05. For isotope labelling: p. value<0.05). For full description of normalization methods, see experimental section.

The significant reduction of the LHCII antenna was clearly observed in the chlorophyll and fluorescence pictures of the native PAGE (Figure 2b). In addition, the amount of PSI in HL was about 60% of that measured in LL thylakoids, when normalized to the amount per cell (Figure 2b, compare lanes 1 and 3). The amount of trimeric and monomeric LHCII complexes was considerably lower in HL as compared to LL thylakoids. A massive accumulation of colored pigments, mostly carotenes and/or xanthophylls (see below), was observed at the leading edge region of the HL sample in the native PAGE and the upper fractions of the sucrose density gradients (Figures 2a and 2b). Analysis of the chlorophyll *a* and *b* in each complex revealed that most of the chlorophyll *b* was located in the PSII-LHCII and LHCII. The chlorophyll *a/b* ratio changed from 2.3 to 5.3 in LL and HL LHCII samples, respectively (Figure 1c). In addition, differences in the carotenoid content were mostly found to be associated with a small fraction of LHCII in the HL thylakoids, appearing to be the major site requiring protection in HL conditions (Figures 2C and S6). Moreover, the enormous increase in the carotenoids/chlorophyll ratio in LHCII, from 0.2 in LL to 1.5 in HL, suggests the participation of additional, still unidentified carotenoid binding sites in LHCII (Figures 2C and S6 and see below). In addition, the enormous accumulation of chlorophyll and carotenoids binding proteins, ELIP and CBR in HL thylakoids possibly add many more carotenoids binding sites (see below).

The reduction in the amount of the LHC proteins was also observed in the fractionation of the protein-chlorophyll complexes in the second dimension by denaturing SDS-PAGE (Figure 2d). Together, these findings indicate that the size of the major LHCII antenna was most strongly reduced in HL cells, resulting in a decreased light harvesting cross-section and a reduction in the amount of damaging light that is directed to the PSII reaction center. Modulation of the LHCII size by light intensity is well documented in high- and low-light adapted plants and algae (Rochaix and Bassi, 2019; Rochaix, 2020; Stefano *et al*., 2014; Negi *et al*., 2020; Erickson *et al*., 2015; Nicol *et al*., 2019). Nevertheless, the drastic and almost complete elimination of the major trimeric LHCII, leaving a “minimal” PSII in HL thylakoids, seems to be a unique phenomenon of *C. ohadii* and its close relative *C. sorokiniana* (not shown).

### HL cells contain reduced amounts of LHC proteins

Proteomics analysis and immunoblotting were performed to analyze protein profiles of HL and LL thylakoids. Since the *C. ohadii* genome sequences were not yet available (NCBI accession PRJNA573576) (Treves *et al*., 2020), the proteomics analysis and protein identification were performed using the genomic nucleotide sequence of its close relative, *C. sorokiniana* (ENA accession LHPG02000000) (Arriola *et al*., 2018). For the immunoblotting analysis, commercial antibodies generated against Arabidopsis LHCII and LHCI proteins were used. In order to identify the *C. ohadii* LHC protein that reacts with a particular antibody, the amino acid sequences of *C. ohadii, C. sorokiniana, Chlamydomonas reinhardtii* and *Arabidopsis thaliana* LHC proteins were aligned and the most reasonable identity of the corresponding antibody determined (Table S3). Using this method, it was possible to identify the two minor-inner LHCII subunits, CP26 and CP29, and to distinguish them from the LHCII major-trimeric and LHCI subunits. However, it was impossible to distinguish between the different major trimeric LHCII subunits and therefore these are grouped together (Figure 2d). Two methods were used to analyze the LC-MS/MS proteomics data: isotope labelling and label-free quantification methods (see Methods and Figure S7). The analyses identified a significant (3-10 fold) reduction in major LHCII subunits content in HL as compared to LL thylakoids and when normalized to the amount per cell, in agreement with the protein complex analysis by the sucrose density gradients and native-PAGE (Figure 2a-c). Similar levels of change were obtained using the immunoblotting and the two proteomic methods (Figures 2d and 3a).

**Figure 3.**
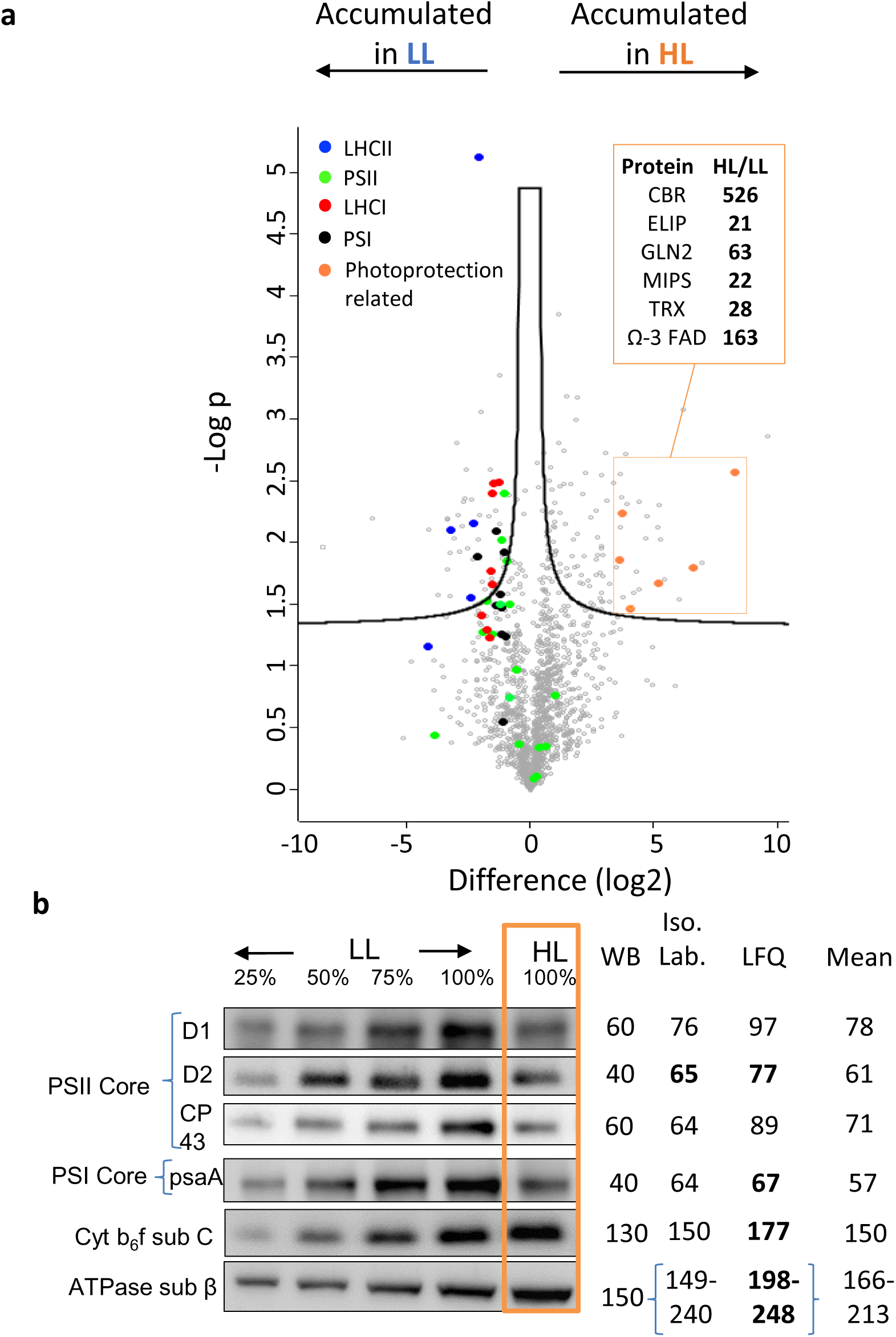
Substantial accumulation of photo-protection proteins and differential expression of photosynthesis proteins in HL and LL thylakoids. **(a)** Mass spectrometry (MS) volcano plot, based on label-free quantification (LFQ) results, showing accumulation of protein in LL (left side) and HL (right side) thylakoids, <u>normalized for equal protein amount of HL and LL</u>. Proteins of interest are colored, as indicated. Significance determined with a permutation-based FDR rate of 0.05. The enormous accumulation of several photo-protection proteins in HL thylakoids is listed to the right, displaying their relative accumulation in HL, as compared to LL cells. This was done by division of the amounts in LL by 1.7, since there are 1.7 more proteins in HL as compared to LL thylakoids (Table 1). The presented results are significant with FDR<0.05. For a full list of the relevant detected proteins, see Table S2. **CBR** – carotene biosynthesis-related, **ELIP** – early light-induced protein, **GLN2** – glutamine synthetase 2, **MIPS** – mio-inositol phosphate synthase, **TRX** – thioredoxin, Ω-3 **FAD** – Ω-3 fatty acid desaturase. (**b**) Immunoblotting was performed to quantify subunits of PSII, PSI, cytochrome *b_6_f* and ATP synthase in HL and LL thylakoids (left side). Values in the table indicate the relative abundance of the proteins in HL compared to LL thylakoids, as measured via western blot (WB), isotope labelling mass spectrometry (Iso. Lab.) and label-free quantification (LFQ) mass spectroscopy (right side). Bold numbers indicate statistically significant changes (for LFQ: FDR<0.05; for Iso. Lab: p. value<0.05). Since there are several proteins annotated as the chloroplast β-subunit of ATPase in the proteome of *C. sorokiniana*, the range of the values obtained is presented. Results in panel b were normalized to demonstrate the <u>change per cell</u>, as described above and in the Experimental section. For full proteins names and details, see Table S2.

The concentrations of the inner PSII antenna proteins, CP26 and CP29, were about 40-50% lower than those measured in LL thylakoids, when normalized per cell (Figures 2d and 3. Table S2). These results correlated well with changes in the amounts of the PSII-RC proteins, D1 and D2, which were about 60-70% lower than those measured in LL thylakoids (Figure 3b). Taken together, PSII retained its inner antennae, i.e., CP26 and CP29, in HL thylakoids. In contrast, the quantities of the major LHCII antenna proteins, LHCB2, B4 and B6, were drastically reduced to about 10-30% of the amounts in LL thylakoids (Figures 2d and 3 and Table S2). Interestingly, the LHCSR proteins that were shown to play an important role in protection against PI in the green alga *Chlamydomonas reinhardtii* (Girolomoni *et al*., 2019; Pinnola, 2019; Vecchi *et al*., 2020; Erickson *et al*., 2015; Rochaix, 2020; De La Cruz Valbuena *et al*., 2019), are absent in *C. ohadii* and *C. sorokiniana.* This may have been the result of their evolutionary elimination since in the absence of the trimeric LHCII, the LHCSR does not protect from PI. Notably, the LHCSR genes are present in another *Chlorella* species, *C. vulgaris* (Cecchin *et al*., 2019).

PsbS is an additional stress-related protein that maybe contribute to the significant NPQ observed in HL cells. Two genes encoding *psbS* homologues are present in the genomes of *C. sorokiniana* and *C. ohadii*. Our proteomics and immunologic analyses of the HL and LL thylakoids did not identify any psbS protein. However, *psbS* and *psbS2* transcripts, which were increased under HL treatment, were identified in a transcriptome analysis of HL and LL cells in a different study (Treves *et al*., 2020). In addition, it has been previously found that the expression of psbS protein in *C. sorokiniana* cells is restricted to autotrophic growth, and was not detected when the alga was grown on an acetate-containing medium (Cecchin *et al*., 2018). Therefore, further study is needed to determine the conditions under which the psbS protein accumulates in HL thylakoids of *C. ohadii* and its possible contribution to the marked NPQ.

A similar analysis of the 10 LHCI subunits was performed, and revealed that the relative amounts of 9 of the LHCI subunits in HL thylakoids were reduced to approximately 50% of the amount, similar to the changes in the PSI-RC protein profile and what was observed by native PAGE, proteomics and immunoblotting analyses (Figures 2 and 3 and Table S2). Together, these observations suggest remarkable HL-triggered modulation of the amounts of proteins of the major trimeric LHCII antenna, which collects and tunnels light to the PI-sensitive PSII. PSI, on the other hand, which is not the main target of PI, is only minimally affected in its protein contents by HL.

### Substantial accumulation of photo-protective proteins in thylakoids of HL-exposed cells

Two methods of LC-MS/MS were employed to quantify differences in protein accumulation in thylakoids of HL and LL cells. As described in the previous section, the most significantly down-regulated proteins in HL thylakoids were the major trimeric LHCII components. LHCI proteins were reduced in HL thylakoids to the same extent (30-40%) as PSII and PSI reaction centers and inner antenna proteins, as was the photosynthetic activity of the linear electron flow (Figure 3 and Table 1). In contrast to these groups of proteins, most of the other thylakoid proteins detected by the MS techniques accumulated in HL thylakoids, explaining the observation that HL thylakoids contain 1.7-fold more protein per cell, when compared to LL thylakoids. Similarly, the subunits of cytochrome *b_6_f* and of the ATP synthase were upregulated by about 160% in HL as compared to LL thylakoids (Figure 3b and Table S2), suggesting increased activity of the PSI cyclic electron flow pathway. The oxygen radical scavenging proteins, the carotenoid biosynthesis enzymes and several enzymes of the PSII repair cycle (Li *et al*., 2019; Liu *et al*., 2019; Chotewutmontri *et al*., 2020) accumulated in HL thylakoids to about 150-200% of the LL levels, when normalized per cell (Figures 3 and 5. Table S2).

Importantly, a group of photoprotection-related proteins markedly accumulated (20-500-fold) in HL as compared to LL thylakoids. These included LHC-like proteins, such as the early light inducible protein (ELIP), and the carotenoid biosynthesis-related protein (CBR). The two are very similar proteins, both shown to bind free chl *a* (but not chl *b*) and xanthophylls (lutein and zeaxanthin) (Figure 3 and Table S2) (Adamska *et al*., 1999; Triantaphylidès and Havaux, 2009; Cecchin *et al*., 2018; Montané *et al*., 1998). These findings suggest the massive activation of a photoprotection mechanism of the accumulation of ELIP and CBR which bind chlorophylls that are in close vicinity to xanthophylls, enabling efficient-energy dissipation of the excited chlorophylls, thereby preventing ROS formation (Son *et al*., 2019). Another important observation was the dramatic increase in the amount of Ω-3 fatty acid desaturase (Ω-3 FAD) in HL thylakoids. It has been suggested that thylakoid membranes rich in polyunsaturated fatty acids (PUFA) serve as structural antioxidants by capturing ROS, thereby protecting the photosynthetic apparatus proteins (Schmid-Siegert *et al*., 2016; Mène-Saffrané *et al*., 2009). Another photoprotection and photosystem repair-related protein that was upregulated was thioredoxin, a redox modulator protein known to participate in stress protection reactions (Liu *et al*., 2019; Lu, 2016; Treves *et al*., 2020). The amount of thioredoxin detected in association with HL thylakoids was 30-fold that associated with LL thylakoids, when normalized per cell (Figure 3 and Table S2). For a full list of additional relevant proteins and their relative quantification, see Table S2.

### Photo-protective carotenoids accumulate in HL cells

Carotenoid accumulation and the xanthophyll cycle are well known PI prevention mechanisms, driven by reactive radical scavenging and sequestering (Nishiyama and Murata, 2014; Niyogi, 1999; Erickson *et al*., 2015). To determine whether carotenoids accumulate in response to HL, as suggested by the yellowish color of the HL cells/thylakoids and by the accumulation of associated enzymes, HL and LL thylakoids were analyzed for their pigment content using high-performance liquid chromatography with a photo diode array detector (HPLC-PDA). The analysis identified a 1.3-1.8-fold accumulation of lutein, neoxanthin and β-carotene in HL thylakoids, when normalized per cell (Figure 4). In addition, antheraxanthin and zeaxanthin, two xanthophylls that are components of the photo-protective xanthophyll cycle, were significantly upregulated or exclusively found in HL thylakoids (Figure 4). In contrast, violaxanthin was reduced in HL as compared to LL thylakoids, as expected when the xanthophyll PI protection cycle is activated (Niyogi, 1999; Erickson *et al*., 2015; Leonelli *et al*., 2017; Nishiyama and Murata, 2014; Murata *et al*., 2007). In parallel, a high correlation was found between upregulation of the carotenoid biosynthesis enzymes and the accumulated amounts of carotenoids in HL thylakoids (Figure 5).

**Figure 4:**
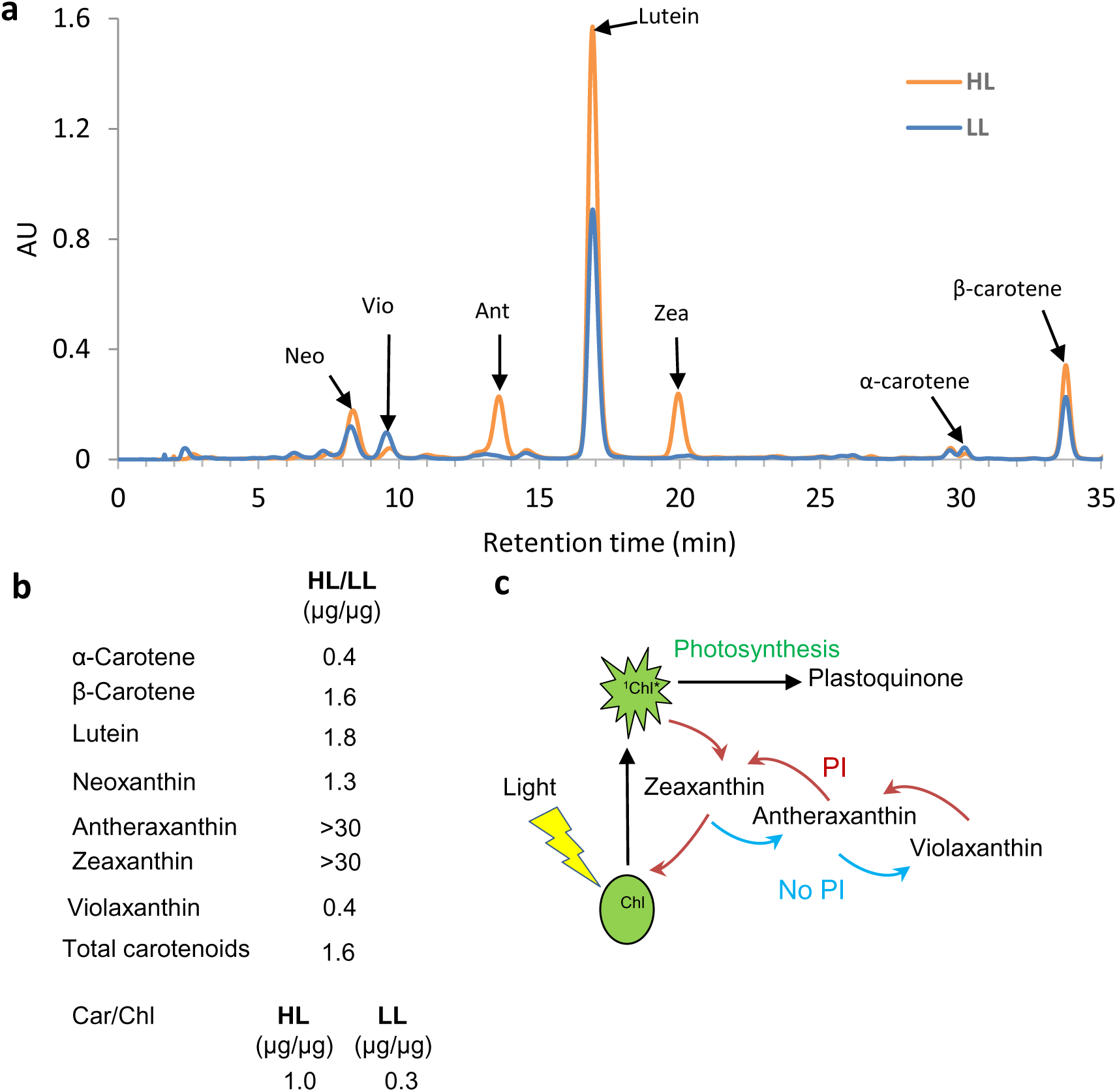
Photoinhibition-protective carotenoids accumulate in HL thylakoids. **(a)** Carotenoids were extracted from LL and HL thylakoids and analyzed by HPLC-PDA separation. To demonstrate changes per cell in the elution profile, HL thylakoid containing 50% the amount of chlorophyll compared to LL samples, were analyzed. Specific carotenoids that were identified using the corresponding markers, are indicated. **b.** The accumulation of carotenoids in HL compared to LL thylakoids as measured by the HPLC-PDA detector. The HL to LL ratio as normalized per cell is shown. See Table S1 for the measured amounts of each carotenoid in μg/cell (x10^-8^). Neo: neoxanthin, Vio: violaxanthin, Ant: antheraxanthin, Zea: zeaxanthin. Results are based on three biological samples of LL and HL thylakoid membranes, harvested on different days. For further details, see Table S1. **c**. The xanthophyll PI protection cycle.

**Figure 5.**
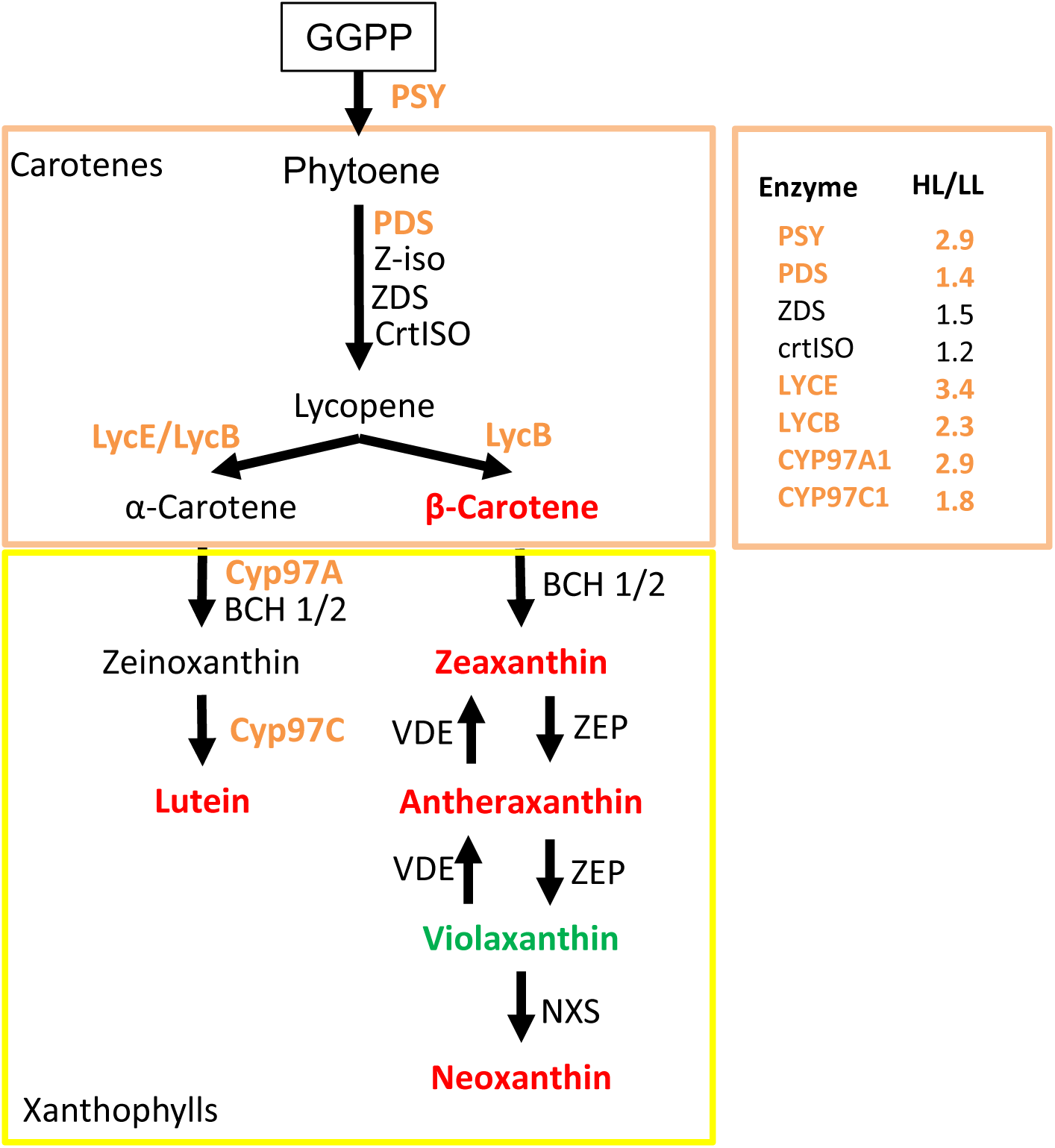
Carotene and xanthophyll biosynthesis enzymes accumulate in HL thylakoids. The carotenoid biosynthesis pathway in chlorophytes is presented, with carotenoids that accumulate in HL thylakoids, colored in red, and the one accumulated in LL thylakoids, colored in green (see Figure 4). Accumulation of enzymes of the carotenoid biosynthesis pathway in HL, as compared to LL cells was determined using the LFQ MS proteomics method. Enzymes showing statistically significant accumulation in HL are colored orange. The amounts detected in HL and LL cells is presented in Table S2 and the accumulation in HL versus LL cells is presented to the right. Bolded and orange-colored enzymes showed statistically significant accumulation (FDR<0.05). GGPP: geranylgeranyl pyrophosphate; PSY: phytoene synthase; PDS: phytoene desaturase; Z-ISO: ζ-carotene isomerase; CrtISO: carotene isomerase; LYCE: lycopene ε -cyclase; LYCB: lycopene β cyclase; CYP97C: carotene ε -monooxygenase; CYP97A: cytochrome P450 hydroxylase; BCH1/2: carotene β-hydroxylase; ZEP: zeaxanthin epoxidase; VDE: violaxanthin de-epoxidase; NXS: neoxanthin synthase.

The carotenoid contents of the different photosynthetic complexes of LL and HL cells were analyzed by extracting the carotenoids of the complexes that were separated by the sucrose density gradients, as shown in Figures 2 and S6, and subjecting them by HPLC. The analysis detected the appearance of the xanthophyll cycle pigments, anteraxanthin and zeoxanthin, in all complexes in HL cells, alongside a massive accumulation of lutein restricted to LHCII and the low-density fraction of the gradient, named here as “free pigments”. β- and α-carotenes accumulated in HL exclusively in the PSI and PSII (Figures 2 and S6). The ∼7-fold increase in lutein levels in HL LHCII, resulted in a lutein to chlorophylls ratio of about 1:1, implying the creation of new carotenoid binding sites in HL LHCII, in additional to the four already known and defined sites (Croce *et al*., 1999). In addition, the massive accumulation of ELIP and CBS in HL thylakoids presumably add many additional binding sites. Neoxanthin was present in all complexes including the PSI-LHCI, in agreement with previous reports of its purification with algal PSI-LHCI and its binding to LHCa5 and LHCa6 in *C. reinhardtii* (Huang *et al*., 2021; Su *et al*., 2019; Qin *et al*., 2015; Pan *et al*., 2021). A cryoEM structural analysis is underway to define the molecular binding sites of the carotenoids in the photosynthetic complexes of HL thylakoids.

Taken together, PI-protecting carotenoids, such as lutein, neoxanthin and β-carotene, accumulated in the corresponding complexes of HL thylakoids, enhancing their ability to dissipate excess light energy. In addition, the known protective xanthophyll cycle was highly active under HL as compared to LL conditions. Therefore, in addition to the significant reduction in the size of the LHCII antenna to protect PSII from PI, a second protective mechanism involving the activation of the xanthophyll cycle and the accumulation of protective carotenoids, operates in HL cells to provide further protection from PI. An additional possibility is that the carotenoid accumulation and the activation of the xanthophyll cycle are short-term protection mechanisms, while the reduced antenna cross-section size is a long-term process (see below). The massive accumulation of carotenoids can also account for the strong quenching of the fluorescence in HL cells (Figure 1).

### The PSII repair cycle is prominently active in HL cells, resulting in rapid turnover of the D1 and D2 PSII proteins

A major PI protection mechanism is the rapid degradation of the damaged PSII proteins, particularly of D1, followed by the synthesis and assembly of newly active PSII (Nishiyama and Murata, 2014; Li *et al*., 2009; Lu, 2016; Keren and Krieger-Liszkay, 2011; Li *et al*., 2018; Cruz *et al*., 2005; Erickson *et al*., 2015; Vecchi *et al*., 2020; Liu *et al*., 2019; Rochaix, 2020; Adir *et al*., 2003; Chotewutmontri and Barkan, 2020). In order to explore how significantly this PI repair process functions under HL conditions, *C. ohadii* cells were incubated under HL and LL, in the presence or absence of chloramphenicol (CAP), which inhibits translation of proteins in the chloroplast, where D1 and D2 are encoded. Thylakoids were isolated 0.5, 1 and 2 h thereafter and the relative amounts of the proteins were determined by immunoblotting using specific antibodies (Figures 6 and S8 and S9). Under HL of 2500 μE m^-2^s^-1^, there was a very rapid turnover of the PSII-RC proteins. The most rapidly degraded protein was D1, displaying a half-life of about 0.5 h, followed by D2, with a half-life of ∼1.0 h; the PSII 43 kD subunit had a significantly longer half-life (Figures 6 and S8, S9). The amounts of D1 and D2 were only slightly reduced in the absence of the antibiotic, indicating rapid translation of the corresponding proteins. Similar results were obtained when a different 70S ribosome inhibitor, lincomycin, was used (not shown). These degradation and translation rates are similar to those reported in cyanobacteria (Mulo *et al*., 2012; Komenda *et al*., 2000; Tyystjärvi *et al*., 1994), but are significantly faster than those reported for the green algae *Chlamydomonas reinhardtii* and for higher plants (Larkum *et al*., 2020; Mulo *et al*., 2012; Liu *et al*., 2019; Rochaix, 2020; Lu, 2016; Schuster *et al*., 1988). These findings differ from a previous report, which showed that D1 in *C. ohadii* was not degraded in cells illuminated by HL (Treves *et al*., 2016). In the PSII repair cycle, D1 is degraded by the FtsH proteases. We indeed were able to identify an accumulation of the members of the FtsH family of proteins in HL thylakoids, which correlated with increased degradation of D1 (Table S2). The other photosynthetic proteins, including the chloroplast-encoded subunits of cyt *b_6_f*, Rubisco and PSI, as well as the nuclear-encoded LHCII, remained relatively stable throughout the HL treatment (Figures 6 and S9). Taken together, these findings imply that *C. ohadii* maintains an active PSII during HL growth by replacing the damaged PSII proteins extremely rapidly, possibly hitting a top record in eukaryotic photosynthetic organisms.

**Figure 6.**
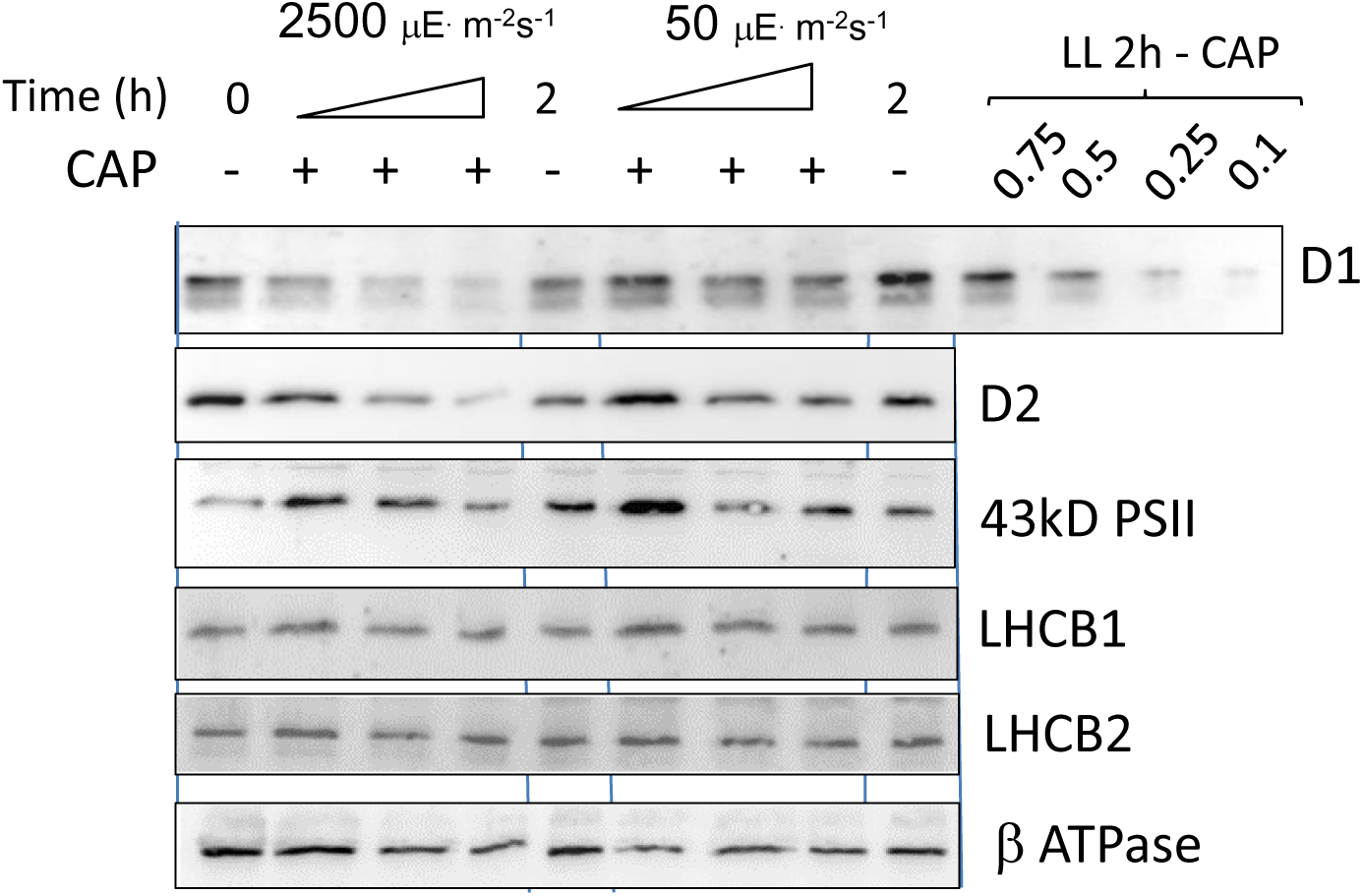
The PSII repair cycle rapidly replaces D1 and D2 in high light. Chloramphenicol (CAP) was added to log-phase *C. ohadii* cells grown under low-light (LL) intensity (50 μE·cm^-2^s^1^). Half of the cell culture was transferred to high light (HL) (2500 μE·cm^-2^s^-1^) and half continued in LL incubation. Samples were collected after 0.5, 1.0 and 2.0 h of incubation for the CAP treated cells, and after 2.0 h of incubation for cells not treated with CAP (Figure 6 and Figures S8 and S9). Thylakoids were isolated and analyzed by SDS-PAGE and immunoblotted using specific antibodies, as indicated. The antibodies raised against the Arabidopsis LHCB1 and LHCB2 recognize one of the *C. ohadii* lhcB2/4/6 proteins (Table S3). The chlorophyll *a/b* ratio did not change and was 5.0 throughout the entire experiment; therefore, thylakoids containing equal amounts of chlorophyll (and proteins) were loaded to each lane of the immunoblot assay.

### Adaptation of *C. ohadii* to LL and HL is a long process

Analysis of the thylakoids of HL and LL *C. ohadii* cells revealed major structural and functional differences that enabled rapid growth under the extreme HL conditions of the desert soil crust. These changes included the upregulation or downregulation of many proteins, chlorophylls and carotenoids and could be the result of modulated gene expression and the involvement of retrograde signal(s). To determine the length of the LL to HL adaptation process, we next explored the timeframe of the transformation between the two structural PSII stages. To this end, cells in the logarithmic growth phase were diluted to 0.2 OD_750_ and then transferred from LL to HL, after which changes in the chl *a/b* ratio were monitored (Figure 7). LL-to-HL adaptation proved to be a long process that took about 8 h. The chl *a/b* ratio remained mostly unchanged during the initial 4-5 h in HL, which is the duplication time of the cells in our system. Thereafter, the chl *a/b* ratio rapidly increased, reaching a value of 10, which essentially reflects almost depletion of chl *b*, after 10 h. This timescale does not align with the daily fluctuations of the light intensities and the life cycle of *C. ohadii* in the desert. During the long summer days in the desert, starting from sunrise, there is about 4 h of relatively low light intensity and high humidity caused by fog and dew. Then, there are about 10 h of HL intensity, severe drought and heat, followed by a 10 h night period (Treves *et al*., 2020). It is assumed that photosynthesis and growth occur during the initial 4 h of the day, while PI and drought occur throughout the rest of the day. The PSII structural changes of reducing the amount of the trimeric LHCII content measured here (Figure 7) do not follow this daily cycle and may involve a longer adaptation process, perhaps changing according to the seasonal fluctuation of light intensity in the desert.

**Figure 7.**
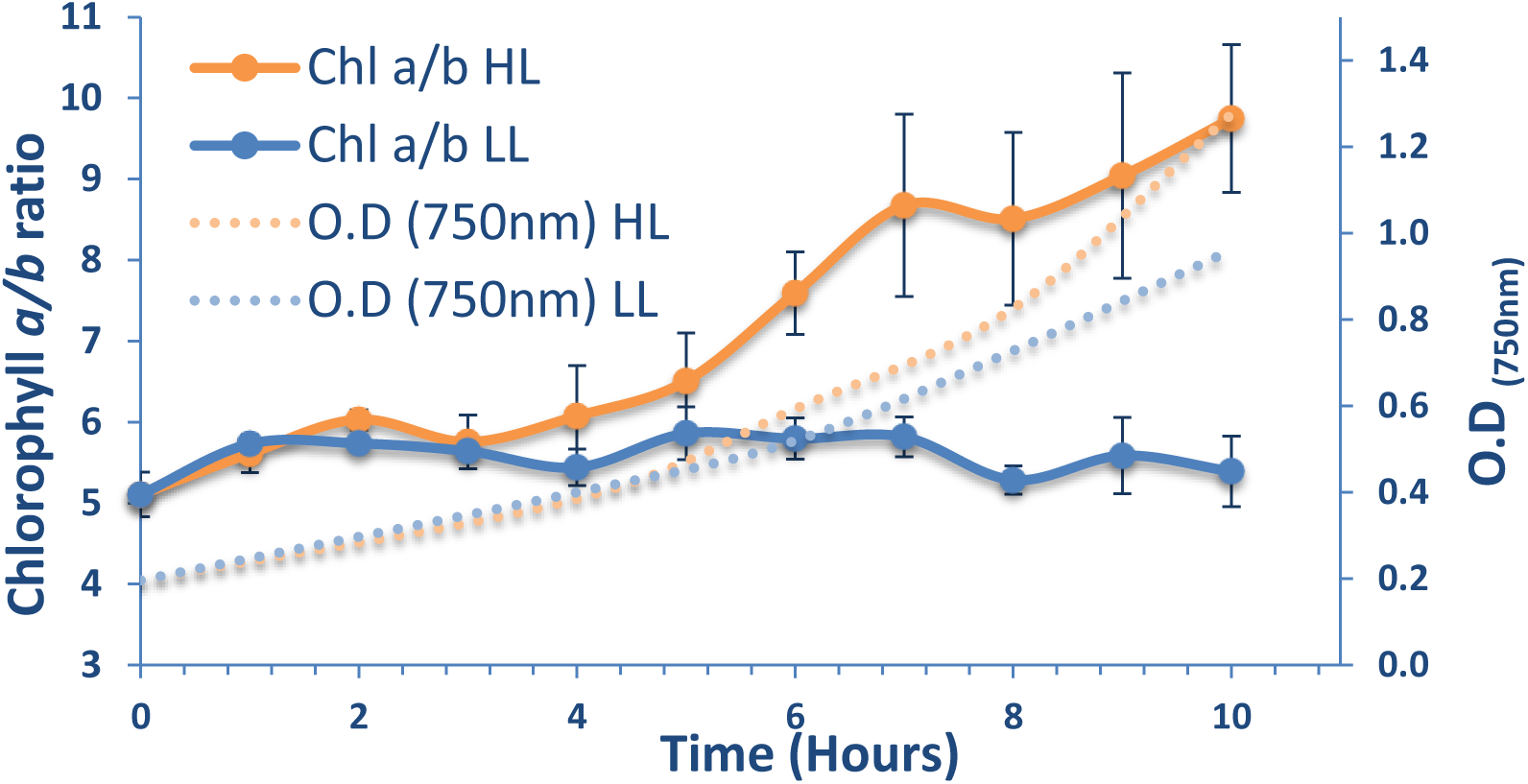
Transition of *C. ohadii* between LL and HL states is a long term process. Log phase-grown LL cells were maintained in LL conditions or transferred to HL. Samples were withdrawn from the culture at the times indicated and the levels of chl *a* and *b* were spectroscopically determined (filled lines). The multiplication rates of the cells were determined by the absorption at 750 nm (dashed lines).

## Conclusions

The desert soil crust is one of the most severe and challenging habitats for photosynthesis organisms. These organisms are bombarded by extreme high light intensities each day, accompanied with high temperatures and severe drought in the summer. The organisms living in such a habitat, mostly cyanobacteria and some green algae, must have developed unique adaptation mechanisms. Alternatively, the evolutionarily developed protection mechanisms that are present in most organisms could have been developed further in order to achieve maximal protection and enable growth in the desert soil crust. The identification and isolation of the green algae *C. ohadii* enabled us to study the molecular mechanisms allowing it to thrive and which protect it from PI at this harmful environment. Our analysis of the thylakoids of cells adapted for several generations to an extreme HL intensity revealed a massive enhancement of the three already known photoinhibition protection mechanisms, exhibiting perhaps the highest recorded activities in plants and algae. Accordingly, no unique properties or new mechanisms that were not identified before in other photosynthetic organisms were noted. Although fluctuations of the PSII antenna size is a well-documented mechanism for adaptation to different light intensities in plants and algae, *C. ohadii* take it to the extreme by reducing the amount of nearly all the trimeric LHCII (Figure 8). In this manner, the specific activity of PSII is increased and most importantly, less harmful ROS are produced within PSII. Nevertheless, the elimination of the peripheral LHCII in HL is a long-term process and cannot fluctuate in the unique diel cycle of desert soil crusts, as may be the case with the massive accumulation of photo-protective proteins in HL thylakoids that was detected in the proteomic analysis. This adaptation process is likely to be the result of significant changes in the modulation of gene expression by the nuclear and the chloroplast genomes and may be facilitated by retrograde signals between the chloroplast and the nucleus. The other two PI-protection processes, carotenoid accumulation and activation of the xanthophyll protection cycle and the extensive operation of the PSII repair cycle to replace the reaction center proteins faster than other plants and algae, can occur on short time scales, thereby facilitating a rapid PSII response to HL. In conclusion, our observations revealed that *C. ohadii* can thrive under HL in the harsh habitat of the desert soil crust by expanding the capacities of the evolutionarily evolved PI protection mechanisms to top-record performance.

**Figure 8.**
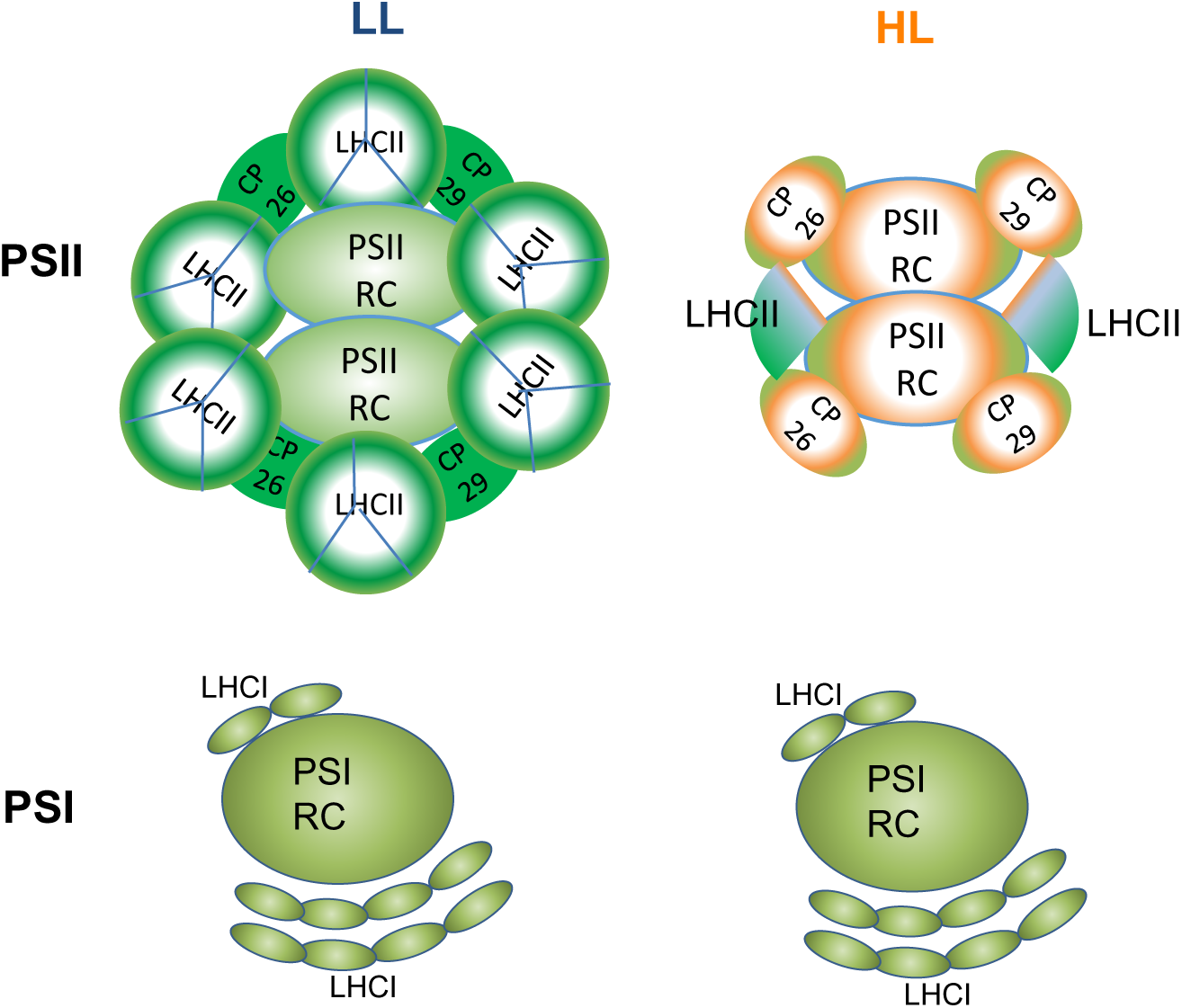
Schematic drawing of PSI and PSII in LL and HL cells. The PSII is schematically presented in its dimeric form. The amount of the trimeric LHCII is significantly reduced in HL cells. The protein structure of PSI does not change between LL and HL cells. The orange color in HL complexes represents the accumulation of carotenoids. The PSI structure is presented here according to the PSI structure of *Chlamydomonas reinhardtii* (Suga *et al*., 2019).

## Methods

### Culture conditions

The isolation of *Chlorella ohadii* from desert soil crust was previously described and the strain was kindly provided by Aaron Kaplan (Hebrew University of Jerusalem) (Treves *et al*., 2013). *C. ohadii* was grown in Tris-acetate-phosphate (TAP) medium at 28°C. Cultures were grown in Erlenmeyer flasks under white light of either 2500 (HL) or 50 μE m^-2^s^-1^ (LL), provided by several LED or fluorescent lamps. To achieve full adaptation to HL, the cells were grown in HL for 6 generations and kept below OD_750nm_ of 0.7 by repetitive dilutions, to prevent self-shading.

### Cells counting

Cells were counted using an Invitrogen™ Countess II Automated Cell Counter. <u>Transmission electron microscopy</u>: Samples were centrifuged (6,000 *g*, 5 min), washed shortly with MES buffer and resuspended in 1 ml fixative (2% paraformaldehyde and 2% glutaraldehyde in 0.1 M cacodylate buffer, pH 7.4) for 90 min at room temperature (RT). Cells were then centrifuged again and embedded in 3% agarose. Thereafter, pellets were incubated in 1 ml secondary fixative (1% OsO_4_ in 0.1 M cacodylate buffer and 1.5% potassium ferrocyanide) for 1 h, on ice. This was followed by incubation of the samples in increasing concentrations of ethanol for dehydration, after which, they were embedded in Epon 812 and baked at 60 °C, for 48 h. Sections (75 nm) were cut using an ultramicrotome (LEICA EM UC7) equipped with a diamond knife, and then transferred to 200 mesh copper grids. The thin sections were analysed using a Zeiss UltraPlus FEG -scanning electron microscope equipped with STEM detector, at an accelerating voltage of 30 keV.

### Fluorescence induction kinetic measurements

Equal numbers of LL or HL cells were dark-adapted for 30 min and the fluorescence induction traces were analyzed using a FL-3500 (Photon Systems Instruments, Brno, Czech Republic) fluorimeter using the manufacturer’s protocols. For determination of PSII antenna size, cells were then illuminated with low light in the presence of 100 μM DCMU. The fluorescence traces were normalized to Fm and the F_0_ of non-DCMU-treated samples was extrapolated, as described in (Tian *et al*., 2019).

### Fluorescence emission spectrum at 77K

Equal numbers of LL and HL cells were dark-adapted for 1h and then loaded into glass tubes, which were placed in a clear glass chamber filled with liquid nitrogen. Excitation was performed at 430 nm and the 445-800 nm emission spectrum was measured using the Fluorolog fluorimeter (Horiba Scientific, Edison, NJ, USA).

### Chlorophyll concentration

Cells were pelleted (max speed, 10 min) followed by pigment extraction in 100% methanol for 10 min, with vigorous vortexing. Cell debris was removed by centrifugation and absorbance was measured at 652 nm and 665 nm. Chlorophyll concentration was determined using the formulas listed in Porra et al. (Porra *et al*., 1989). To determine chlorophyll concentrations in isolated thylakoids, the same procedure was performed using 80% acetone instead of methanol, and accordingly, absorbance was measured at 645 nm and 663 nm (Porra *et al*., 1989).

### Protein quantification

Thylakoid membranes were solubilized in 1% β-dodecyl maltoside (DDM) for 20 min at RT, and then cleared by centrifugation (15,000 rpm, 10 min). Protein concentration was then determined using the Bradford assay (Biorad), with bovine serum albumin (BSA) serving as a standard.

### Large-scale thylakoid preparation

Log phase-grown cells (1-6 L) were pelleted (4500 rpm, 2 min), washed once and resuspended in cold STN1 buffer (0.4 M sucrose, 10 mM tricine pH 8.0, 10 mM NaCl), at 4°C. Cells were broken by passing them through a microfluidizer operating at 50 psi; unbroken cells removed by low-speed centrifugation (2000 RPM, 15 min, 4°C). The thylakoids were then pelleted by high-speed centrifugation (2 h, 50,000 rpm, 4°C), resuspended in a minimal volume of cold STN1 and stored in aliquots at −80°C.

### Small-scale thylakoid preparation

Log phase-grown cells (50-100 ml, OD_750_<0.7) were pelleted (4500 rpm, 2 min), resuspended in 0.5 ml TMNS buffer (50 mM Tris pH=8.0, 5 mM MgCl_2_, 10 mM NaCl and 0.4 M sorbitol) and disrupted by 3 cycles of 30-sec sonication, in ice water, performed using a micro-tip sonicator. Unbroken cells were removed by centrifugation (1000 rpm, 2 min). After pelleting the thylakoids (10,000 rpm, 10 min, 4°C), they were re-suspended in a minimal volume of TMNS buffer.

### Protein immunoblotting

Thylakoid proteins were fractionated by denaturing SDS-PAGE, and then blotted onto nitrocellulose membranes. The membranes were stained with Ponceaou S, blocked and then incubated overnight with antibodies recognizing different photosynthesis proteins (Agrisera). Following incubation with the second antibody (peroxidase-conjugated goat-anti-rabbit) for 1 h, signals were detected by chemoluminescence using the Fusion 400 instrument (Halpert *et al*., 2019). The amino acid sequences of the LHC proteins of *C. ohadii, C. sorokiana, C. renhardtii* and *A. thaliana* were aligned using the pBlast tool, to predict which antibody raised against the Arabidopsis LHC subunit interacts with the corresponding *C. ohadii* LHC subunit, as listed in Table S3. Antibodies were purchased from Agrisera. Product numbers are as following: lhcb1 - AS01 004, lhcb2 - AS01 003, lhcb4 - AS04 045, lhcb5 - AS01 009, lhcb6 - AS01 010, lhca2 - AS01 006, psbA - AS11 1786, psbC - AS11 1787, psbD - AS06 146, psaA - AS06 172, petC - AS08 330.

### Native PAGE electrophoresis

Thylakoid membranes were solubilized for 20 min, at 25°C, in 2% β-dodecyl maltoside (DDM), 0.7% n-octyl-β-d-glucoside (OG) and 1% sodium dodecyl sulfate (SDS), and then cleared by centrifugation (12,000 *g*, 10 min). Thereafter, the extracted protein complexes were fractionated on a gradient 3-12% acrylamide-PAGE that was run at 70V until the front line reached the bottom of the gel. Gels were photographed to enable observation of the chlorophyll protein complexes. In addition, the gel was analysed for the fluorescence of the fractionated complexes, measured with a *Typhoon FLA 7000*, at an excitation wavelength of 532 nm. For the second-dimension fractionation in SDS-PAGE, the native-PAGE lane was soaked in denaturing SDS-PAGE sample buffer, heated to 85°C for 5 min and loaded on a 12% SDS-PAGE. Proteins were detected by Coomassie blue R or silver staining.

### Sucrose density gradient separation of chlorophyll protein complexes

Thylakoid membranes were solubilized for 10 min, at 4°C, in 1% α-DDM and then cleared by centrifugation (12,000 *g*, 10 min). Thereafter, the extracted protein complexes were fractionated in a sucrose density gradient of 4-45% in 25 mM MES (pH 6.5) buffer, by centrifugation, at 100,000 *g*, for 20 h, using SW-41 rotor. The different protein chlorophyll complexes were separately collected and further analysed for spectroscopic, chlorophyll, carotenoid and protein content.

### PSII activity measurements

#### DCPIP reduction

Thylakoids (10 or 20 μg chlorophyll) were suspended in buffer A (0.4 M sucrose, 50 mM MES, pH 6, 10 mM MgCl_2_, 5 mM CaCl_2_) in the presence or absence of 1 mM of the electron donor 1-5-diphenylcarbazide (DPC). The sample was then incubated for 40 sec in the dark, with 50 or 75 μM 2,6-dichlorophenolindophenol (DCPIP) before absorbance was measured at 595 nm. The sample was then illuminated with white light (1000 μE m^-2^s^-1^) for 20 sec and absorbance was measured again at 595 nm. Specific PSII activities were 171 (± 22) and 260 (±40) µmols DCPIP mg chl^-1^ h^-1^ for LL and HL thylakoids, respectively.

#### O_2_ evolution

Thylakoids containing 10 μg chlorophyll were incubated in buffer A, supplemented with 150 mM 2, 6-dichloro-1, 4-benzoquinone (DCBQ), for 3 min, in the dark, before turning on the light and measuring the maximum rate of O_2_ evolution using a Clark-type electrode. Specific PSII activities were 225 (± 26) and 328 (±63) µmol O_2_ mg chl^-1^ h^-1^ for LL and HL thylakoids, respectively.

#### Sample preparation for mass-spectroscopy analysis

Four thylakoid samples from different biological repeats (two of each HL and LL thylakoids) were re-suspended in 0.1 M HEPES, 1% sodium deoxycholate (SDC), pH 8.0, and incubated for 10 min, at 95°C. Protein concentration was determined using the Bradford/BCA assay. Protein (100 μg) from each sample was reduced with 5 mM DTT, for 30 min, at 65°C, and then alkylated with 12.5 mM iodoacetamide (IAA), for 30 min, at RT, in the dark. Excess IAA was quenched with 10 mM DTT, and then digested overnight, at 37 °C, with sequencing-grade trypsin (Promega), added at a ratio of 1:100 (trypsin to total protein, w/w). Each sample was split into two aliquots. For **label-free analysis**, phase transfer was performed to remove sodium deoxycholate (Masuda *et al*., 2008). For the **reductive dimethylation method**, LL and HL samples were incubated, for 30 min, at 37°C, in 20 mM light (^12^CH_2_O) or heavy (^13^CD_2_O) formaldehyde and 40 mM cyanoborohydride. The reaction was terminated and quenched by the addition of 100 mM glycine. The heavy- and light-labeled samples were then mixed and subjected to phase transfer (Masuda *et al*., 2008). The resulting peptides were desalted using a double-layered C_18_ Stage tip (Rappsilber *et al*., 2007).

#### LC−MS/MS

Desalted peptides were analyzed by liquid chromatography coupled mass spectrometry using a Q Exactive Plus mass spectrometer (Thermo Fisher Scientific), coupled online to a RSLC nano HPLC (Ultimate 3000, UHPLC Thermo Fisher Scientific). The peptides were resolved by reverse-phase chromatography on 0.075 × 180 mm fused silica capillaries (J&W) packed with Reprosil reversed-phase material (Dr. Maisch; GmbH, Germany). The peptides were eluted with a linear 60 min gradient of 5–28%, followed by a 15 min gradient of 28–95%, and a 10 min wash with 95% acetonitrile mixed with 0.1% formic acid in water (at flow rates of 0.15 μl/min). Mass spectrometry analysis was performed in positive mode, at a range of m/z 300– 1800, resolution 60,000 for MS1 and 15,000 for MS2, using a repetitively full MS scan, followed by high-energy collisional dissociation (HCD) of the 10 most dominant ions selected from the first MS scan.

#### Data analysis of the proteomics

Raw data were processed via MaxQuant software (1.6.3.4)(Masuda *et al*., 2008) to obtain label-free quantification (LFQ), and quantification of dimethylated samples. Database searching was performed using trypsin as the digesting enzyme with the following parameters: cysteine carbamidomethylation as a fixed modification, methionine oxidation and N-terminal acetylation as variable modifications, up to two missed cleavages permitted, mass tolerance of 20 ppm, 1% protein false discovery rate (FDR) for protein and peptide identification, and a minimum of two peptides for pairwise comparison in each protein. As the proteome of *C. ohadii* is yet unavailable, the *Chlorella sorokiniana* proteome database, downloaded from Uniprot (ENA accession LHPG02000000), was used for protein identification (20/10/2019, 10,201 sequences).

Statistical analysis and data quality checks were performed using the Perseus software (1.6.2.1) (Tyanova *et al*., 2016). After removing the reversed and known contaminating proteins, the LFQ values or dimethylation heavy/light ratios were log_2_-transformed and the reproducibility across the biological replicates was evaluated. As LL thylakoids contain 1.7-times less protein than HL thylakoids per cell (Table 1), intensities of all proteins in LL samples were divided by 1.7 to normalize the results per cell.

**For LFQ**, only proteins that were detected in both repeats of at least one group (HL or LL) were included in the analysis. Missing values were replaced by imputation. A two-sample t-test (FDR = 5%) was performed to identify the proteins with significantly changed expression levels. **For reductive dimethylation**, only proteins that were detected in both biological repeats were subjected to a one-sample t-test (p. val= 5%). The initial “row” proteomics DMT and LFQ tables of results are presented in the supplemental data.

#### Carotenoid analysis

Frozen thylakoid samples were weighed and extracted in 8 ml hexane:acetone:ethanol (50:25:25, v/v), followed by 5 min saponification in 80% (w/v in H_2_O) KOH. In some experiments, the saponification step was omitted (e.g. Figure S6). The saponified material was extracted twice with hexane. For the HPLC analysis, the hexane extractions were evaporated in a Savant SpeedVac apparatus (Savant, Holbrook, NY) and suspended in 400 µl methanol:acetone (2:1), passed through a 0.2- µm PTFE filter (Pall) and stored at RT, in the dark, for no more than 24 h before analysis.

High-performance liquid chromatography (HPLC) was performed using a Waters HPLC system equipped with a Waters HPLC 600 pump, a Waters 996 Photodiode Array detector and a 717 plus autosampler. Separation, based on previously described protocol (Rodrigo *et al*., 2013), was carried out with a C30 column (YMCA, 5 µm, 4.6×250 mm) coupled to a C30 guard column system (YMCA) as described (Wei *et al*., 2020). The data were recorded at 250-600 nm and analyzed by the ‘Empower’ software (Waters). Carotenoids were identified by their absorption spectra, retention times and co-injection spiking of authentic standards (Sigma, Carotenature). Quantification was performed based on β carotene calibration curves prepared with an authentic standard (Sigma Aldrich). Carotenoid content was expressed as µg/g frozen weight. Since HL thylakoids contain 50% less chlorophyll per cell than LL thylakoids, 400 µg chlorophyll of LL samples were compared to 200 µg chlorophyll of HL samples.

#### Thylakoid protein turnover

Log-phase cells, grown under LL, were diluted to OD_750_ = 0.4, divided to four and transferred to HL or LL. Chloramphenicol (CAP) at a final concentration of 200 μg/ml was then added to one LL and one HL sample. Lower concentrations of CAP did not inhibit the translation of organelle proteins since the drug did not enter the cells. Cells (50 ml) were collected at the indicated time points and thylakoids were isolated and analysed by immunoblotting. The chl *a/b* ratio did not change during the incubation and, therefore, equal amounts of chl were loaded in each lane.

## Acknowledgments

Funding: Funding was provided by a “Nevet” grant from the Grand Technion Energy Program (GTEP) and a Technion VPR Berman Grant for Energy Research.

We thank Aaron Kaplan for providing *C. ohadii* and *C. sorokiniana* cells, Alex Brandis for carotenoid analysis, Avihai Danon for antibodies, Lihi Shaulov for the electron microscopy analysis, Rachel Nechushtai, Aaron Kaplan and Nir Keren for help and valuable advice and discussions and Rachel Nechushtai for methods and assistance with the experiments. We thank Oded Kleifeld for his invaluable help with the proteomic methods and data analysis. We also thank Rawad Hanna, Sivan Arber, Yitav Gil-Ad, Yasmin Shibli, Yael Herskovitz-Pollak, Zuchman Rina, Roi Siegelman and the Smoler Proteomics Center in the Technion for their help with the proteomics sample preparation and analysis.

## Author contribution

GL and GS designed the research. GL, SK and GS performed the research. BE, VL helped with some experiments. AM, TI and YT performed the carotenoids analysis. GL, GS and NA wrote the article.

## Conflict of interests

The authors declare no competing financial interests.

## Data availability statement

All original data and material information requests can be made to either Gadi Schuster at or Guy Levin at.

## Supporting information

Additional Supporting Information can be found in the online version of this article.

**Figure S1. HL and LL cells have a similar multiplication rate.** *C. ohadii* cells were grown at light intensities of 2500 (HL; red) and 50 mE (LL; blue) m^-2^ s^-1^, to log phase. The cells were then diluted to OD_750_ of 0.2 and continue to grow at the same light intensities. The mean blot and standard error of three independent experiments are shown. The generation time is indicated.

**Figure S2. HL and LL cells have similar amounts of thylakoids.** The lengths of thylakoids in 0.5 μM^2^ of electron-micrographs, as shown in the figure of the HL cell, were determined using the Image J software in HL (red, left) and LL (blue, right) cells.

**Figure S3. HL and LL cells have similar amounts of stacking thylakoids.** Electron-micrographs of HL (red, left) and LL (blue, right) cells were analyzed using the Image J software, in order to determine the amount of stacking of the thylakoids.

**Figure S4. SDS-PAGE of spinach (Spn), *C. ohadii* LL thylakoids (LL) and *C. ohadii* HL thylakoids (HL).** Spinach (Spn) and *C. ohadii* LL thylakoids (1 μg chlorophyll of each), and *C. ohadii* HL thylakoids (0.5 μg chlorophyll) were separated by SDS-PAGE. Note that the HL lane contain 1.7 times the amount of protein, as indicated in Table 1 and Table S1. However, HL thylakoids contain reduced amounts of LHC proteins.

**Figure S5. PSII antenna is reduced in HL cells. a.** 77K fluorescence spectra of HL and LL cells. The emission spectra were normalized to the 723 nm peak of the LHCI and the baseline fluorescence. **b**. The light-limited, maximal PSII electron transport rate in LL and HL cells was determined using subsaturating light using the chlorophyll fluorescence induction curves of cells measured in the presence of DCMU as displayed in Figure 1b. Fitting parameters are indicated in the figure. **b**. The area above the fluorescence induction curves taken at different low light intensities, as shown in Figure 1b, were plotted and the linear fit forced through the zero point. The difference in the slopes indicates 17% functional PSII antenna size in HL, as compared to LL cells

**Figure S6. Carotenoids content in the photosynthetic complexes of LL and HL thylakoids included accumulation of antheraxanthin and zeaxanthin in all HL complexes and massive accumulation of lutein in LHCII.** Carotenoids were extracted from the photosynthetic complexes that were separated by sucrose density gradient as shown in **panel a**, and analyzed by HPLC-PDA separation. Equal amounts of chlorophyll of each LL and HL complex and the “free pigment” zone of the gradient were analyzed. **Panels b-e**: data obtained with the saponification of the samples. **Panels f-i:** data obtained without saponification of the samples. Neo, neoxanthin. Vio, violaxanthin. Ant, antheraxanthin. Zea, zeaxanthin. Chl a, chlorophyll *a*. Chl b, chlorophyll *b*.

**Figure S7. Proteomics analysis of HL and LL thylakoids - data quality check for LFQ MS experiment. a.** Heat map of accumulated proteins in LL (two experiments) and HL (two experiments) thylakoids. Equal amounts of proteins were analysed for each sample. Red indicates accumulation and green indicates depletion. **b.** Scatter plot of identified proteins. X and Y axis demonstrate LFQ intensity in the indicated sample. Pearson correlation value between samples is indicated in blue at the top left corner of each box. **c.** Principal component analysis of LL and HL samples. **d.** Peptide distribution in LL and HL thylakoids. The X axis shows the LFQ intensity and Y axis show the peptide count. Red columns represent proteins undetected by MS and their missing values added based on normal distribution. Data analysis was performed and figures were generated with Perseus software (1.6.2.1). Note that normalization in these analyses were performed on equal amounts of proteins and not the amount per cell.

**Figure S8. Design and SDS-PAGE of the experiment analyzing the turnover of PSII proteins (presented in** Fig. 6 **of the manuscript).** Chloramphenicol (CAP) was added to *C. ohadii* log-phase cells grown under low light intensity of 50 mE m^-2^s^-1^. Half of the cell culture was transferred to HL (2500 mE m^-2^s^-1^) and half continued in LL incubation. Samples were collected following 0.5, 1.0 and 2.0 h of incubation, for the CAP-treated cells, and after 2.0 h, for cells incubated without CAP. Thylakoids were isolated and analyzed by SDS-PAGE and immunoblotting. An example of a stained SDS-PAGE is shown here and the immunoblots in Fig. 6. The numbers of the lanes in red color indicate the time points in the experiment, which is also indicated in the upper panel. The chlorophyll *a/b* ratio at the starting and end of the experiment is indicated. It remained stable at 5.0 throughout the entire experiment. The OD_750_ of the cell cultures are indicated. Thylakoid sample volumes containing equal amounts of chlorophyll (and proteins) were loaded in each lane of the immunoblot assay.

**Figure S9. Turnover of thylakoid proteins in *C. ohadii* in high light (HL) and low light (LL).** The experiments were performed as described in Fig 6 and Extended Data Fig S8. The ponceau S staining of a gel is shown at the bottom.

## Supporting Tables

**Table S1. Differences and similarities of HL versus LL grown cells.** Absorbance, chl, carotenoids, thylakoid proteins and PSII activity were measured and are presented as the content per cell, for an easy comparison. ND-not detected. Yellow background indicates a major difference between the HL and LL. Values are means of at least 3 independent experiments and significance was determined using two-tailed distribution students t test. **+** = p<0.05, **++** =p<0.005, **+++** =p<0.0005.

**Table S2. Differential accumulation of thylakoid proteins in HL versus LL grown cells, as normalized to the amount per cell.** Thylakoid proteins of HL and LL grown cells were analyzed using the LFQ and isotope labelling methods. The amounts in HL versus LL, following normalization to the amount per cell, is presented. This was done by the division of the LL intensities by 1.7, since there are 1.7-fold more proteins in HL as compared to LL thylakoids (Table 1). A full list of relevant proteins detected by MS analysis of HL and LL thylakoids is listed. The annotations were determined via analysis of MS raw data in MaxQuant software with the proteome of *Chlorella sorokiniana*. Values in the “protein” column were manually determined using NCBI and UNIPROT blast tool, based on the sequence of the identified protein and comparison to other photosynthetic organisms. Values highlighted with green background are statistically significant. Statistical significance for LFQ was determined with a permutation-based FDR<0.05. Significance for stable isotope labelling was set as p<0.05. Yellow background indicates a significant up-expression in HL. Pale blue background indicates significant under-accumulation in HL. **ND** – Not detected. Results are based on two biological samples of thylakoid membranes from LL and HL cells, harvested on different days. *Two α and two β sub-units of chloroplastic ATP synthase proteins were identified. The presented values are of the proteins with the HL/LL ratio most similar to the values of the other subunits. The HL/LL ratios of the proteins not included in the table: α-ATP synthase – 1.96 (LFQ), not detected in isotope labelling. β-ATP synthase – 1.93 (LFQ), 1.5 (isotope labelling).

**Table S3. Identification of the C. ohadii LHC proteins and cross-reactivity with antibodies generated against the Arabidopsis LHC proteins.**

**I. Amino acid sequences of “*C. ohadii* LHC proteins”, detected by mass spectrometry, are almost identical to *C. sorokiniana* LHC proteins.** Since the *C. ohadii* proteome was not yet available during this experiment, we used *Chlorella sorokiniana* proteome for protein identification in our mass spectrometry (MS) analysis.

13 proteins were detected by MS and annotated as “Chlorophyll a/b binding protein”. NCBI blastp was used to detect proteins with highest degree of homology to the amino acid (aa) sequences of *C. ohadii* LHC proteins, obtained from Aaron Kaplan. Each sequence of *C. ohadii* LHC subunit displayed the highest identity to one of the LHCs detected by MS, as well as to a subunit of *C. sorokiniana*, allowing us to define the Uniport ID and to further compare it to *C. reihardtii*, in which the identity of each LHC subunit is known (next table).

Five proteins are shown as an example. “MS peptides compared to *C. ohadii*” column indicates the number of peptides of a specific protein with an aa sequence that is also present in the corresponding protein in *C. ohadii* and *C. sorokiniana*. The degree fold of expression in HL as compared to LL, as normalized per cell and determined by the LFQ proteomic method, is listed in the right-most column.

**II. Comparison of the identified LHC subunits to those of *Chlamydomonas reinhardtii* enabled functional identification of the LHC subunits.** *C. reinhardtii* LHC subunits are well annotated. Therefore, to annotate the 13 *C. ohadii* LHC subunits, their aa sequences were aligned to those of *C. reinhardtii.* Out of the 13 highest ranking proteins, 5 were determined as LHCII subunits (**left table**) and 8 were determined as LHCI subunits (**right table**). Note that ranking determined by NCBI blastp depends on the identity as well as other parameters. Each subunit identity (green) was determined based on the highest ranked match. For example, A0A2PU146 = LHCBM4.

**III. Determination of the LHC subunit that reacts with antibodies generated against the *Arabidopsis* LHC proteins.** Immunoblotting (IB) was performed with antibodies (ABs) raised against *Arabidopsis thaliana* LHC (AS01 011, Agrisera). Since *A. thaliana* contains a different number of LHC subunits than *C. ohadii*, it was necessary to determine which of the 13 LHC proteins detected in *C. ohadii* are the most likely to react with each of the ABs. The aa sequences of the *Arabidopsis* proteins, used to raise the Abs, were aligned against the 13 sequences of LHC of *C. ohadii*. For the internal LHC subunits, CP29 and CP26 ABs, the identifications were conclusive as their molecular weights (MWs) matched those of the bands detected in IB (∼30kd), alongside their high sequence homology. For the trimeric LHCII ABs, we could not identify the specific reacting subunit, as their MWs are similar (∼25kd) and both crossreacting ABs (B1 and B2) showed highest similarity to the same subunits. However, there was a slight difference in the MW of the detected bands, indicating that different subunits react with the each of the two different ABs. Blast analysis suggested that B6 and A2 ABs reacted with subunits of LHCI. However, both ABs showed highest similarity to the same subunits, which have similar MWs, making it impossible to distinguish between them. No cross reactivity was achieved for B3, A1, and A3-4 ABs.

**Tables S4-S6:** The “raw” proteomics data of the DMT and LFQ results.

## Notes

### Competing Interest Statement

The authors have declared no competing interest.

