## Supplementary material for "The desert green algae *Chlorella ohadii* thrives at excessively high light intensities by exceptionally enhancing the mechanisms that protect photosynthesis from photoinhibition": Figures S1-S9

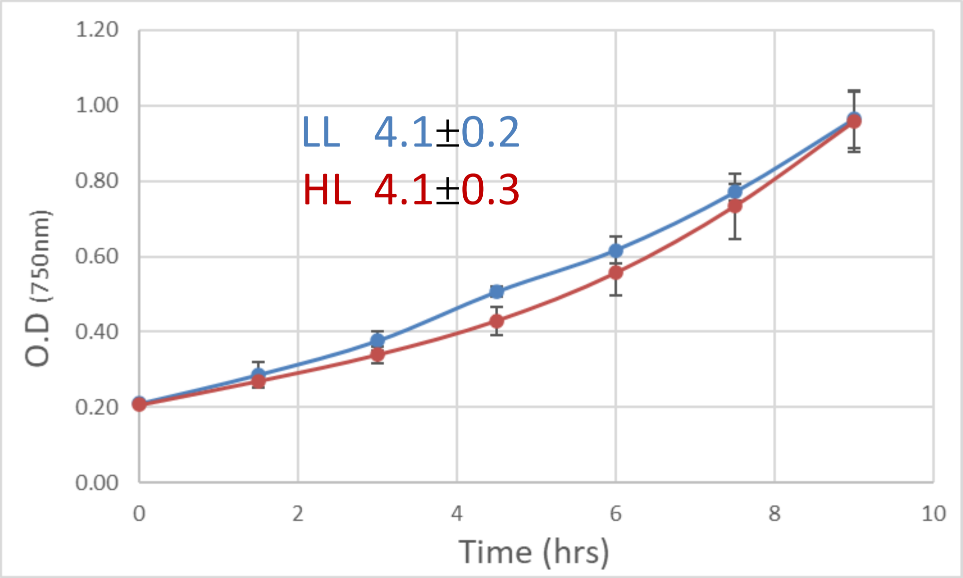


**Figure 8. Schematic drawing of PSI and PSII in LL and HL cells.** The PSII is schematically presented in its dimeric form. The amount of the trimeric LHCII is significantly reduced in HL cells. The protein structure of PSI does not change between LL and HL cells. The orange color in HL complexes represents the accumulation of carotenoids. The PSI structure is presented here according to the PSI structure of *Chlamydomonas reinhardtii* (Suga *et al.*, 2019).

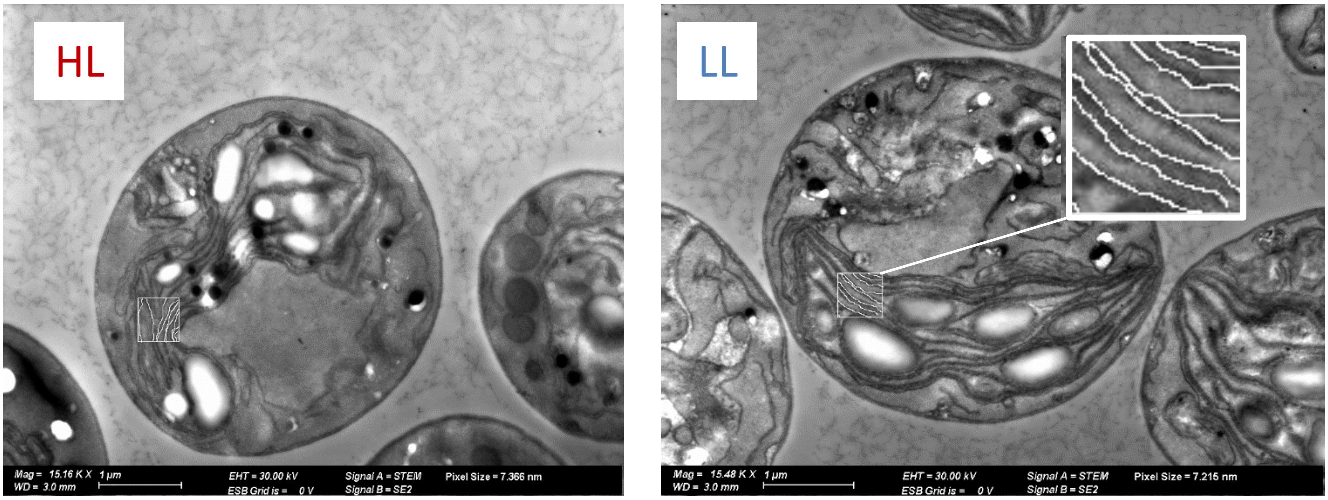


Thylakoids length in 0.5 $\mu M^2$

HL 4.0 ± 0.4 μM
LL 3.6 ± 0.3 μM

**Figure S2. HL and LL cells have similar amounts of thylakoids.** The lengths of thylakoids in 0.5 $\mu M^2$ of electron-micrographs, as shown in the figure of the HL cell, were determined using the Image J software in HL (red, left) and LL (blue, right) cells.

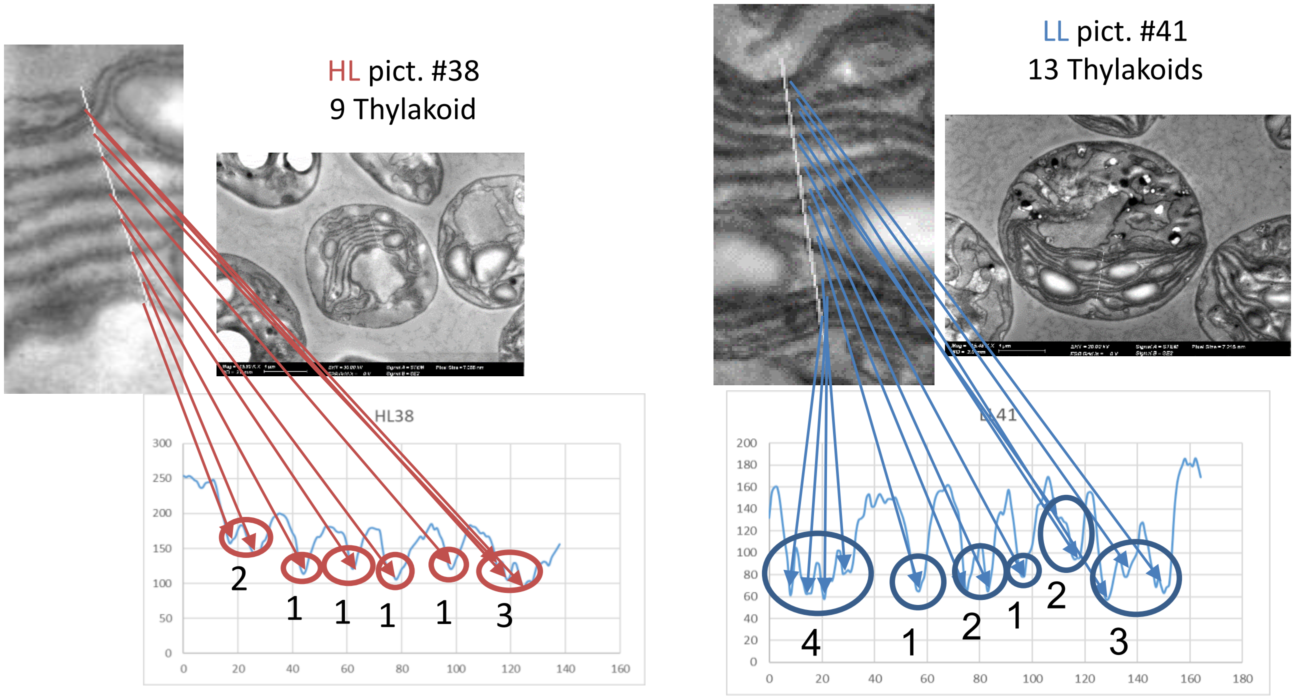


Stacking of thylakoids:
Average LL = 2.0 (±1)
Average HL = 1.7 (±0.7)

**Figure S3. HL and LL cells have similar amounts of stacking thylakoids.** Electron-micrographs of HL (red, left) and LL (blue, right) cells were analyzed using the Image J software, in order to determine the amount of stacking of the thylakoids.

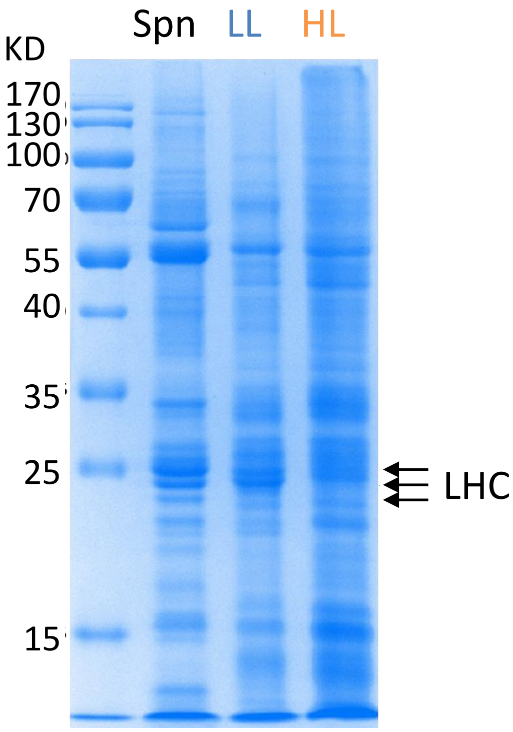


**Figure S4. SDS-PAGE of spinach (Spn), *C. ohadii* LL thylakoids (LL) and *C. ohadii* HL thylakoids (HL).** Spinach (Spn) and *C. ohadii* LL thylakoids (1 µg chlorophyll of each), and *C. ohadii* HL thylakoids (0.5 µg chlorophyll) were separated by SDS-PAGE. Note that the HL lane contain 1.7 times the amount of protein, as indicated in Table 1 and Table S1. However, HL thylakoids contain reduced amounts of LHC proteins.

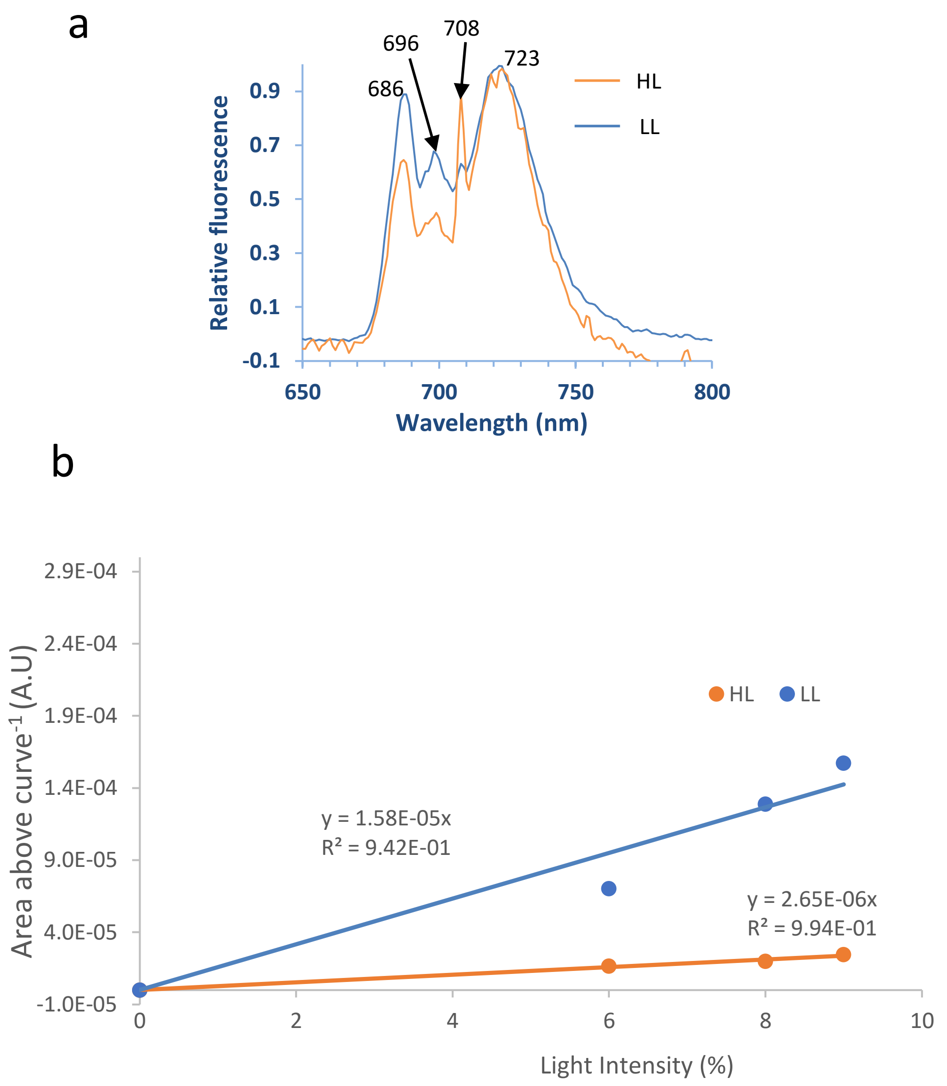


**Figure S5. PSII antenna is reduced in HL cells. a.** 77K fluorescence spectra of HL and LL cells. The emission spectra were normalized to the 723 nm peak of the LHCI and the baseline fluorescence. **b**. The light-limited, maximal PSII electron transport rate in LL and HL cells was determined using subsaturating light using the chlorophyll fluorescence induction curves of cells measured in the presence of DCMU as displayed in Figure 1b. Fitting parameters are indicated in the figure. **b**. The area above the fluorescence induction curves taken at different low light intensities, as shown in Figure 1b, were plotted and the linear fit forced through the zero point. The difference in the slopes indicates 17% functional PSII antenna size in HL, as compared to LL cells

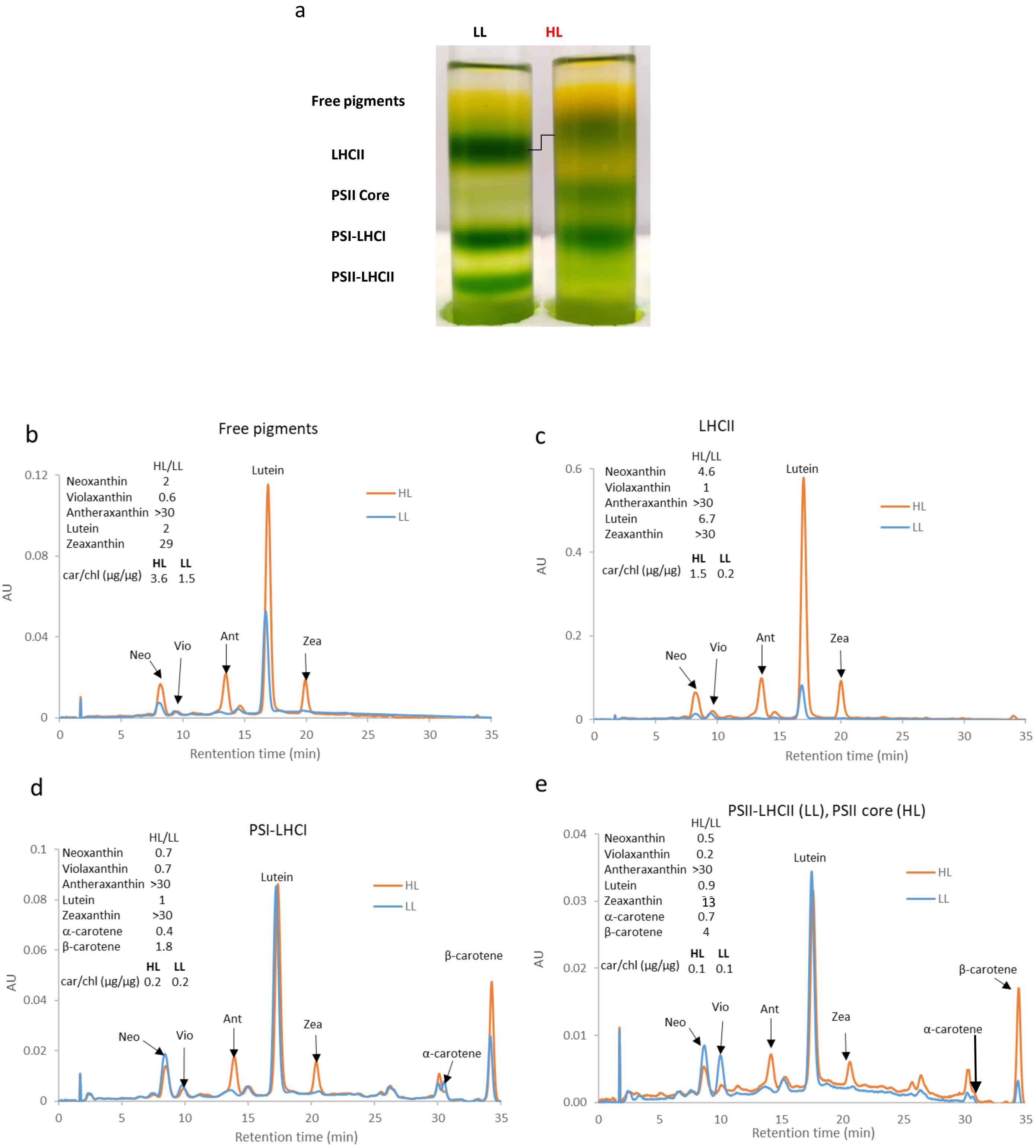

a
LL
HL
Free pigments
LHCII
PSII Core
PSI-LHCI
PSII-LHCII
b
Free pigments
HL/LL
Neoxanthin 2
Violaxanthin 0.6
Antheraxanthin >30
Lutein 2
Zeaxanthin 29
HL LL
car/chl (μg/μg) 3.6 1.5
Neo
Vio
Ant
Lutein
Zea
HL
LL
AU
0.12
0.08
0.04
0
0 5 10 15 20 25 30 35
Rentention time (min)
c
LHCII
HL/LL
Neoxanthin 4.6
Violaxanthin 1
Antheraxanthin >30
Lutein 6.7
Zeaxanthin >30
HL LL
car/chl (μg/μg) 1.5 0.2
Neo
Vio
Ant
Lutein
Zea
HL
LL
AU
0.6
0.4
0.2
0
0 5 10 15 20 25 30 35
Retention time (min)
d
PSI-LHCI
HL/LL
Neoxanthin 0.7
Violaxanthin 0.7
Antheraxanthin >30
Lutein 1
Zeaxanthin >30
α-carotene 0.4
β-carotene 1.8
HL LL
car/chl (μg/μg) 0.2 0.2
Neo
Vio
Ant
Lutein
Zea
α-carotene
β-carotene
HL
LL
AU
0.1
0.08
0.06
0.04
0.02
0
0 5 10 15 20 25 30 35
Retention time (min)
e
PSII-LHCII (LL), PSII core (HL)
HL/LL
Neoxanthin 0.5
Violaxanthin 0.2
Antheraxanthin >30
Lutein 0.9
Zeaxanthin 13
α-carotene 0.7
β-carotene 4
HL LL
car/chl (μg/μg) 0.1 0.1
Neo
Vio
Ant
Lutein
Zea
α-carotene
β-carotene
HL
LL
AU
0.04
0.03
0.02
0.01
0.00
0 5 10 15 20 25 30 35
Retention time (min)

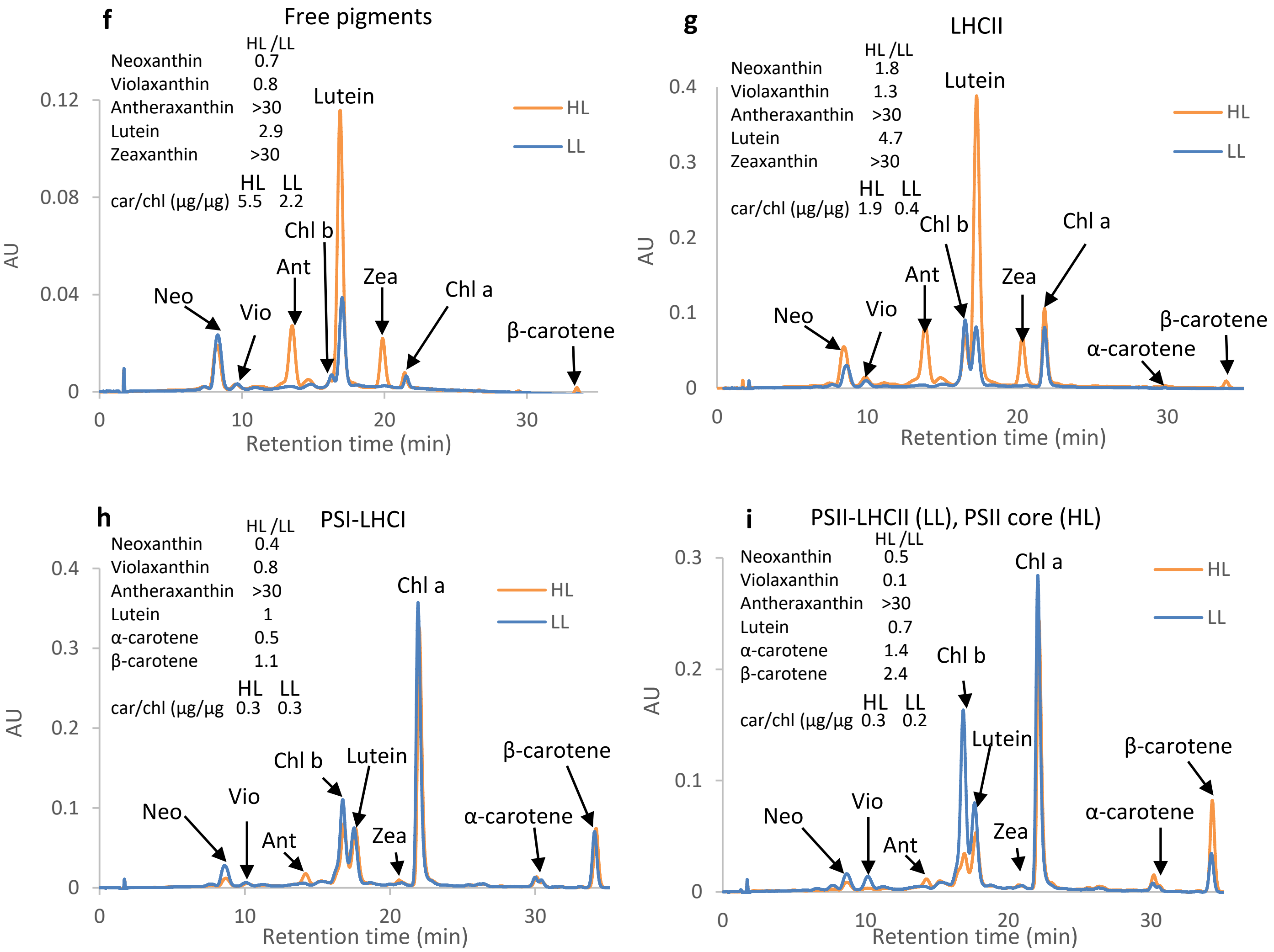


**Figure S6. Carotenoids content in the photosynthetic complexes of LL and HL thylakoids included accumulation of antheraxanthin and zeaxanthin in all HL complexes and massive accumulation of lutein in LHCII.** Carotenoids were extracted from the photosynthetic complexes that were separated by sucrose density gradient as shown in **panel a,** and analyzed by HPLC-PDA separation. **Panels b-e**: data obtained with the saponification of the samples. **Panels f-i:** data obtained without saponification of the samples. Equal amounts of chlorophyll of each LL and HL complex and the "free pigment" zone of the gradient were analyzed. Neo, neoxanthin. Vio, violaxanthin. Ant, antheraxanthin. Zea, zeaxanthin. Chl a, chlorophyll *a*. Chl b, chlorophyll *b*.

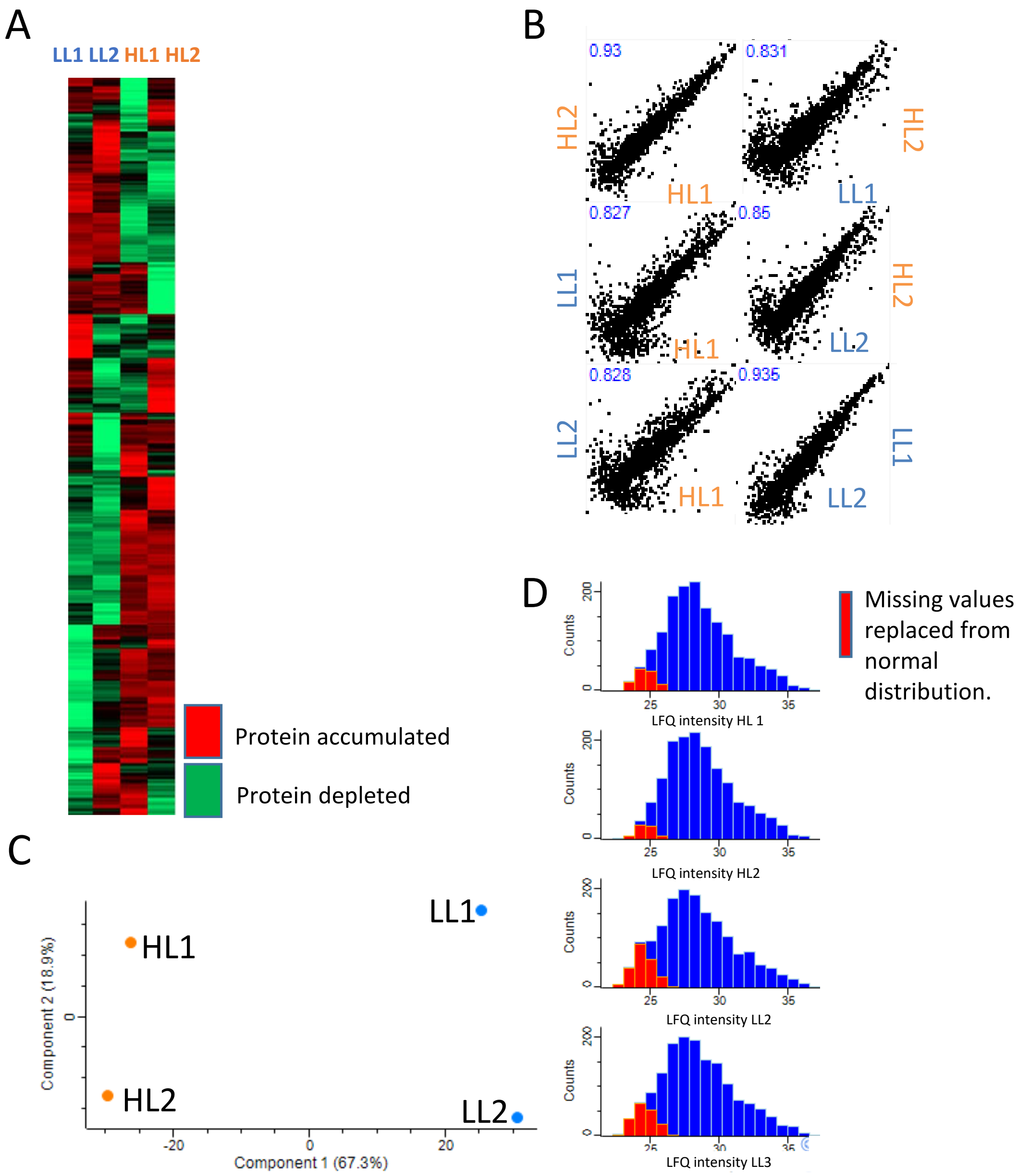


**Figure S7. Proteomics analysis of HL and LL thylakoids - data quality check for LFQ MS experiment. a.** Heat map of accumulated proteins in LL (two experiments) and HL (two experiments) thylakoids. Equal amounts of proteins were analysed for each sample. Red indicates accumulation and green indicates depletion. **b.** Scatter plot of identified proteins. X and Y axis demonstrate LFQ intensity in the indicated sample. Pearson correlation value between samples is indicated in blue at the top left corner of each box. **c.** Principal component analysis of LL and HL samples. **d.** Peptide distribution in LL and HL thylakoids. The X axis shows the LFQ intensity and Y axis show the peptide count. Red columns represent proteins undetected by MS and their missing values added based on normal distribution. Data analysis was performed and figures were generated with Perseus software (1.6.2.1). Note that normalization in these analyses were performed on equal amounts of proteins and not the amount per cell.

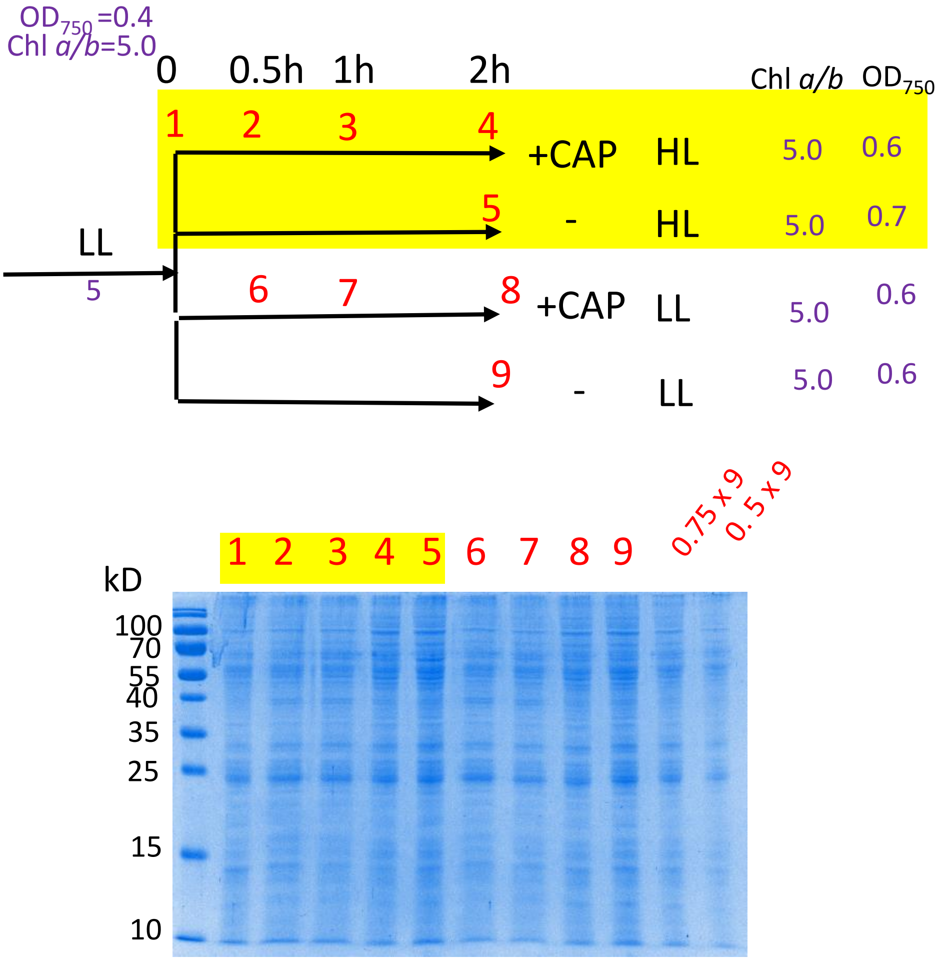


**Figure S8. Design and SDS-PAGE of the experiment analyzing the turnover of PSII proteins (presented in Fig. 6 of the manuscript).** Chloramphenicol (CAP) was added to *C. ohadii* log-phase cells grown under low light intensity of 50 mE $m^{-2}s^{-1}$. Half of the cell culture was transferred to HL (2500 mE $m^{-2}s^{-1}$) and half continued in LL incubation. Samples were collected following 0.5, 1.0 and 2.0 h of incubation, for the CAP-treated cells, and after 2.0 h, for cells incubated without CAP. Thylakoids were isolated and analyzed by SDS-PAGE and immunoblotting. An example of a stained SDS-PAGE is shown here and the immunoblots in Fig. 6. The numbers of the lanes in red color indicate the time points in the experiment, which is also indicated in the upper panel. The chlorophyll *a/b* ratio at the starting and end of the experiment is indicated. It remained stable at 5.0 throughout the entire experiment. The $OD_{750}$ of the cell cultures are indicated. Thylakoid sample volumes containing equal amounts of chlorophyll (and proteins) were loaded in each lane of the immunoblot assay.

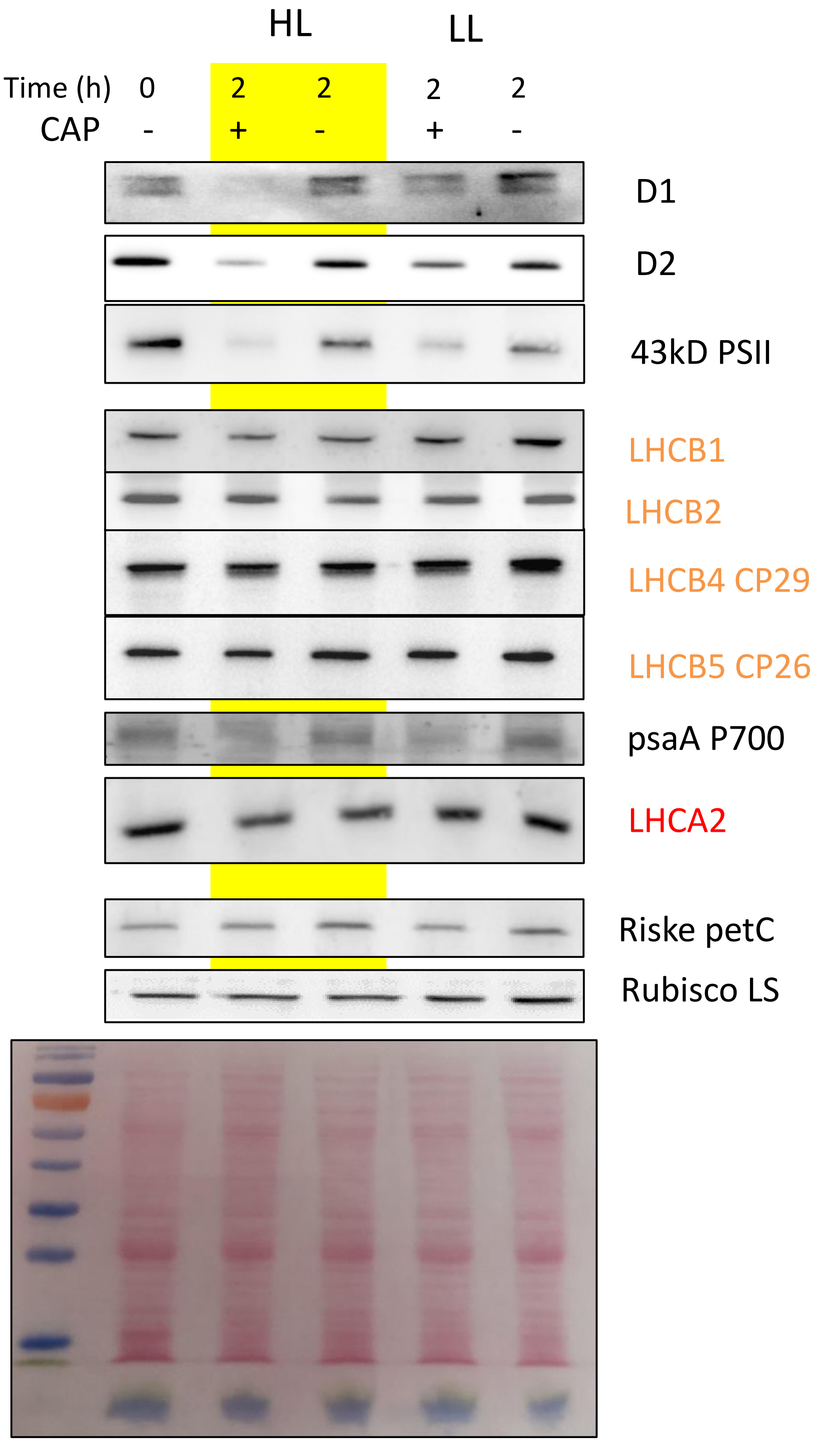


**Figure S9. Turnover of thylakoid proteins in *C. ohadii* in high light (HL) and low light (LL).** The experiments were performed as described in Fig 6 and Extended Data Fig S8. The ponceau S staining of a gel is shown at the bottom.
