## Supplementary material for "The desert green algae *Chlorella ohadii* thrives at excessively high light intensities by exceptionally enhancing the mechanisms that protect photosynthesis from photoinhibition": Tables S1-S2

**Table S1. Differences and similarities of HL and LL grown cells**

| **Parameter** | **HL** | **LL** | **HL/LL** | **Significant** |
| --- | --- | --- | --- | --- |
| Cell size (diameter) μm | 3.7 (±0.7) | 3.9 (±0.5) | 0.95 |  |
| Multiplication time (cell division) hours | 4.1 (±0.2) | 4.1 (±0.3) | 1 |  |
| Thylakoids amount (μm /0.5μm^2^) | 3.6 (±0.3) | 4.0 (±0.3) | 0.9 |  |
| Thylakoids stacking degree | 2 (±1) | 1.7 (±0.7) | 1.2 |  |
| OD_750nm_/Cell ( **X10^-8^** ) | 9.0 | 8.2 | 1 |  |
| Chl *a* μg/Cell ( **X10^-8^** ) | 17 | 32 | 0.5 | +++ |
| Chl *b* μg/Cell ( **X10^-8^** ) | 1.5 | 5.6 | 0.4 | ++ |
| Chl *a/b* ratio | 8.8 | 3.9 |  | ++ |
| Chl μg/Cell ( **X10^-8^** ) | 19 | 38 | 0.5 | +++ |
| Chl μg/OD_750nm_ | 1.4 | 3.4 | 0.4 | +++ |
| Thylakoid Proteins mg/Cell ( **X10^-6^** ) | 3.5 | 2.1 | 1.7 | +++ |
| α-Carotene μg/Cell ( **X10^-8^** ) | 0.11 | 0.29 | 0.4 |  |
| β-Carotene μg/Cell ( **X10^-8^** ) | 1.9 | 1.2 | 1.6 |  |
| Lutein μg/Cell ( **X10^-8^** ) | 10 | 5.8 | 1.8 | + |
| Neoxanthin μg /Cell ( **X10^-8^** ) | 1.3 | 0.99 | 1.3 | + |
| Violaxanthin μg/Cell ( **X10^-8^** ) | 0.27 | 0.67 | 0.4 | + |
| Antheraxanthin μg/Cell ( **X10^-8^** ) | 1.7 | ND | >30 | + |
| Zeaxanthin μg /Cell ( **X10^-8^** ) | 1.7 | 0.04 | >30 | + |
| Total carotenoids μg/Cell ( **X10^-8^** ) | 19 | 11 | 1.6 | + |
| Carotenoid/Chlorophyll ratio (μg/μg) | 0.98 | 0.3 | 3.3 | ++ |
| DPC-DCPIP PSII activity μmol/Cell *1hr ( **X10^-8^** ) | 4.8 | 6.4 | 0.75 | + |
| H_2_O-DCPIP PSII activity μmol/Cell *1hr ( **X10^-8^** ) | 4.7 | 5.9 | 0.8 | + |
| H_2_O-DCBQ PSII activity μmol O_2_/Cell*1hr( **X10^-8^** ) | 6.1 | 8.4 | 0.73 | ++ |
| Fv/Fm | 0.73 | 0.81 | 0.9 | + |

**Table S1. Differences and similarities of HL versus LL grown cells.** Absorbance, chl, carotenoids, thylakoid proteins and PSII activity were measured and are presented as the content per cell, for an easy comparison. ND-not detected. Yellow background indicates a major difference between the HL and LL. Values are means of at least 3 independent experiments and significance was determined using two-tailed distribution students t test. **+** = p<0.05, **++** =p<0.005, **+++** =p<0.0005.

**Table S2. Differential accumulation of thylakoid proteins in HL versus LL grown cells, as normalized to the amount per cell.**

| **PSII core** | Protein | Annotation | LFQ HL/LL | Isotope labelling HL/LL | UNIPROT I.D |
| --- | --- | --- | --- | --- | --- |
|  | psbA | PSII RC protein D1 | 0.97 | 0.76 | W8SIR2_CHLSO |
|  | psbB | PSII CP47 RC protein | 0.82 | 0.62 | W8TIK4_CHLSO |
|  | psbC | PSII CP43 RC protein | 0.89 | 0.64 | W8SUG2_CHLSO |
|  | psbD | PSII RC D2 protein | 0.77 | 0.65 | W8SYD4_CHLSO |
|  | psbH | PSII RC protein H | >0.01 | 0.70 | A0A2I4S6N1_9CHLO |
|  | psbH | PSII RC protein H (2) | 1.18 | 0.46 | W8SIT0_CHLSO |
|  | psbL | PSII RC protein L | 0.12 | ND | W8SKK5_CHLSO |
| **Cyt. b_559_ (PSII)** | psbE | Cyt. b559 subunit alpha | 0.98 | 0.68 | W8SU91_CHLSO |
|  | psbF | Cyt. b559 subunit beta | 0.75 | 0.62 | W8SIQ3_CHLSO |
| **OEC of PSII** | psbO | 33kDa OE of PSII | 0.45 | 0.51 | A0A2P6TWR3_CHLSO |
|  | psbP | OE enhancer 2 of PSII | 0.60 | 0.41 | A0A2P6U3U0_CHLSO |
|  | psbP | psbP chloroplastic (2) | 2.22 | 1.99 | A0A2P6TKP3_CHLSO |
|  | psbQ | OE enhancer 3 of PSII (isoform A) | 0.52 | 0.42 | A0A2P6TGD3_CHLSO |
| **Damage, assembly, and repair of PSII components** | psbR | PSII 10 kDa | 1.92 | 0.47 | A0A2P6TSU7_CHLSO |
|  | psb28 | PSII RC psb28 protein | 2.68 | 1.90 | A0A2P6TPR3_CHLSO |
|  | psb27 | PSII repair psb27 | 1.26 | ND | A0A2P6TFK0_CHLSO |
|  | LPA | Low PSII accumulation | 1.82 | 1.47 | A0A2P6TD59_CHLSO |
|  | hcf136 | PSII stability assembly factor | 3.42 | 2.85 | A0A2P6U5H8_CHLSO |
|  | Hcf244 | NAD(P)-binding Rossmann-fold superfamily | 1.55 | 1.63 |  |
|  | OHP2 | Low CO2 and stress-induced one-helix | 3.65 | ND |  |
|  | psbN | Protein PsbN | 2.06 | ND | W8SUC6_CHLSO |
|  | ftsh1 | ATP-dependent zinc metalloprotease isoform B | 2.14 | 1.97 | A0A2P6U3P5_CHLSO |
|  | ftsh2 | ATP-dependent zinc metalloprotease FtsH | 1.81 | 1.64 | W8SY61_CHLSO |
|  | ftsh3 | ATP-dependent zinc metalloprotease FtsH mitochondrial-like | 3.66 | ND | A0A2P6TQC9_CHLSO |
|  | EF-Tu | Elongation factor Tu | 1.33 | 1.17 | W8TIL6_CHLSO |
| **PSI** | psaA | PSI P700 chlorophyll a apoprotein A1 | 0.67 | 0.64 | W8SY74_CHLSO |
|  | psaB | PSI P700 chlorophyll a apoprotein A2 | 0.82 | 0.70 | W8SUA3_CHLSO |
|  | psaC | PSI iron-sulfur centre | 0.79 | 0.57 | W8SKM2_CHLSO |
|  | psaD | PSI RC subunit | 0.73 | 0.70 | A0A2P6TKF8_CHLSO |
|  | psaE | PSI RC subunit IV | 0.77 | 0.82 | A0A2P6U4S6_CHLSO |
|  | psaF | PSI RC subunit III | 0.80 | 0.63 | A0A2P6TPV8_CHLSO |
|  | psaH | PSI RC subunit VI | 0.67 | 0.56 | A0A2P6TPU7_CHLSO |
|  | psaK | PSI RC subunit | 0.39 | ND | A0A2P6U0J1_CHLSO |
|  | psaL | PSI RC subunit XI | 0.86 | 0.59 | A0A2P6TC44_CHLSO |
| **PSI assembly** | ycf3 | PSI assembly protein Ycf3 | 3.66 | 3.22 | W8SKR6_CHLSO |
|  | ycf4 | PSI assembly protein Ycf4 | 3.16 | 2.83 | W8SUD8_CHLSO |
| **Cytochrome b6f** | petA | Cytochrome f | 1.45 | 1.23 | W8SIU9_CHLSO |
|  | petB | Cytochrome b_6_ | 2.90 | 1.32 | W8SIS5_CHLSO |
|  | petC | Cytochrome b_6_f complex iron-sulfur | 1.77 | 1.50 | A0A2P6TNS6_CHLSO |
|  | petD | Cytochrome b_6_f complex subunit 4 | 1.18 | 1.26 | W8SUC0_CHLSO |
| **ATP synthase subunits** | atpA | ATPsyn-α* | 2.72 | 2.39 | W8SKS1_CHLSO |
|  | atpB | ATPsyn-β* | 2.48 | 2.39 | W8TIQ2_CHLSO |
|  | atpC | ATPsyn-γ | 2.89 | 2.18 | A0A2P6TTQ2_CHLSO |
|  | atpD | ATPsyn-δ | 2.13 | 1.86 | A0A2P6TN97_CHLSO |
|  | atpE | ATPsyn-ε | 2.04 | 2.25 | W8SYF3_CHLSO |
| **LHCII** | LHCB5-CP29 | Chlorophyll a-b binding | 0.33 | 0.36 | A0A2P6U4D0_CHLSO |
|  | LHCB5-CP26 | Chlorophyll a-b binding | 0.35 | 0.31 | A0A2P6TTG5_CHLSO |
|  | LHCBM2 | Chlorophyll a-b binding | 0.40 | 0.26 | A0A2P6TDA6_CHLSO |
|  | LHCBM4 | Chlorophyll a-b binding | 0.18 | 0.11 | A0A2P6U146_CHLSO |
|  | LHCBM6 | Chlorophyll a-b binding | 0.10 | ND | A0A2P6TNN6_CHLSO |
| **LHCI** | LHCA1 | Chlorophyll a-b binding | 0.55 | 0.52 | A0A2P6TT36_CHLSO |
|  | LHCA2 | Chlorophyll a-b binding | 0.44 | 0.41 | A0A2P6TMX4_CHLSO |
|  | LHCA4 | Chlorophyll a-b binding | 0.62 | 0.45 | A0A2P6TQ14_CHLSO |
|  | LHCA5 | Chlorophyll a-b binding | 0.58 | 0.40 | A0A2P6U4K1_CHLSO |
|  | LHCA6 | Chlorophyll a-b binding | 0.59 | 0.42 | A0A2P6TPR7_CHLSO |
|  | LHCA7 | Chlorophyll a-b binding | 0.60 | 0.55 | A0A2P6TS63_CHLSO |
|  | LHCA8 | Chlorophyll a-b binding | 0.70 | 0.50 | A0A2P6TZ50_CHLSO |
|  | LHCA9 | Chlorophyll a-b binding | 0.50 | 0.41 | A0A2P6TMI2_CHLSO |
| **Carotenoid biosynthesis** | CHYB | Beta-carotene hydroxylase | 0.19 | ND | A0A2P6TMA3_CHLSO |
|  | CrtISO | Carotene isomerase | 1.24 | 1.10 | A0A2P6TG93_CHLSO |
|  | CYP97A1 | Lutein deficient | 2.88 | 2.83 | A0A2P6U1R2_CHLSO |
|  | CYP97C1 | Carotene epsilon-isoform A | 1.84 | 1.41 | A0A2P6TIE5_CHLSO |
|  | DXR | 1-deoxy-D-xylulose 5-phosphate reductoisomerase | 3.55 | ND | A0A2P6TBN0_CHLSO |
|  | DXS | 1-deoxy-D-xylulose 5-phosphate synthase | 2.76 | ND | A0A2P6TD92_CHLSO |
|  | ISPG | 4-hydroxy-3-methylbut-2-en-1-yl diphosphate synthase | 2.61 | ND | A0A2P6TKU4_CHLSO |
|  | ISPH | 4-hydroxy-3-methylbut-2-enyl diphosphate reductase | 1.38 | ND | A0A2P6TLB6_CHLSO |
|  | LYCB | Chloroplast lycopene beta cyclase | 2.25 | ND | A0A2P6TYX4_CHLSO |
|  | LYCE | Lycopene epsilon cyclase | 3.39 | ND | A0A2P6TQM7_CHLSO |
|  | PDS | Phytoene desaturase | 1.42 | 1.16 | A0A2P6TR05_CHLSO |
|  | PSY | Phytoene synthase | 2.90 | ND | A0A2P6TDH2_CHLSO |
|  | ZDS | Zeta-carotene desaturase | 1.47 | 1.29 | A0A2P6TFB0_CHLSO |
| **ROS**  **scavenging** | APX | Thylakoid-bound ascorbate peroxidase | 0.23 | ND | A0A2P6TXN1_CHLSO |
|  | CAT | Catalase | 1.48 | 1.88 | A0A2P6TUJ2_CHLSO |
|  | GPX | Glutathione peroxidase | 1.78 | 1.25 | A0A2P6TI90_CHLSO |
|  | GR | Glutathione reductase | 0.48 | 0.51 | A0A2P6TMT4_CHLSO |
|  | GST | Microsomal glutathione S-transferase 3 | 1.97 | ND | A0A2P6U1D1_CHLSO |
|  | SOD | Superoxide dismutase | 2.14 | ND | A0A2P6TBW4_CHLSO |
| **Photoprotection related (major changes)** | CBR | Carotene biosynthesis related | 526.99 | 24.19 | A0A2P6TNM2_CHLSO |
|  | ELIP | Early light-inducible | 21.18 | ND | A0A2P6U2C2_CHLSO |
|  | TRX | Thioredoxin | 28.00 | ND | A0A2P6TV63_CHLSO |
|  | MIPS | Myo-inositol-1-phosphate synthase | 22.39 | ND | A0A2P6TVK9_CHLSO |
|  | GLN2 | Glutamine synthetase | 63.40 | 5.27 | A0A2P6TID2_CHLSO |
|  | FAD | Omega-3 fatty acid desaturase | 163.57 | 16.22 | A0A2P6U195_CHLSO |

**Table S2. Differential accumulation of thylakoid proteins in HL versus LL grown cells, as normalized to the amount per cell.** Thylakoid proteins of HL and LL grown cells were analyzed using the LFQ and isotope labelling methods. The amounts in HL versus LL, following normalization to the amount per cell, is presented. This was done by the division of the LL intensities by 1.7, since there are 1.7-fold more proteins in HL as compared to LL thylakoids (Table 1). A full list of relevant proteins detected by MS analysis of HL and LL thylakoids is listed. The annotations were determined via analysis of MS raw data in MaxQuant software with the proteome of *Chlorella sorokiniana*. Values in the “protein” column were manually determined using NCBI and UNIPROT blast tool, based on the sequence of the identified protein and comparison to other photosynthetic organisms. Values highlighted with green background are statistically significant. Statistical significance for LFQ was determined with a permutation-based FDR<0.05. Significance for stable isotope labelling was set as p<0.05. Yellow background indicates a significant up-expression in HL. Pale blue background indicates significant under-accumulation in HL. **ND** – Not detected. Results are based on two biological samples of thylakoid membranes from LL and HL cells, harvested on different days. *Two α and two β subunits of chloroplastic ATP synthase proteins were identified. The presented values are of the proteins with the HL/LL ratio most similar to the values of the other subunits. The HL/LL ratios of the proteins not included in the table: α-ATP synthase – 1.96 (LFQ), not detected in isotope labelling. β-ATP synthase – 1.93 (LFQ), 1.5 (isotope labelling).

on different days. *For α and β subunits of chloroplastic ATP synthase there were two identified proteins each. The presented values are of the proteins with the HL/LL value which is more similar to the values of the other subunits. The HL/LL values of the proteins not included in the table: α-ATP synthase – 1.96 (LFQ), not detected in isotope labelling. β-ATP synthase – 1.93 (LFQ), 1.5 (Isotope labelling).
