## Supplementary material for "The desert green algae *Chlorella ohadii* thrives at excessively high light intensities by exceptionally enhancing the mechanisms that protect photosynthesis from photoinhibition": Table S3

### Slide 1
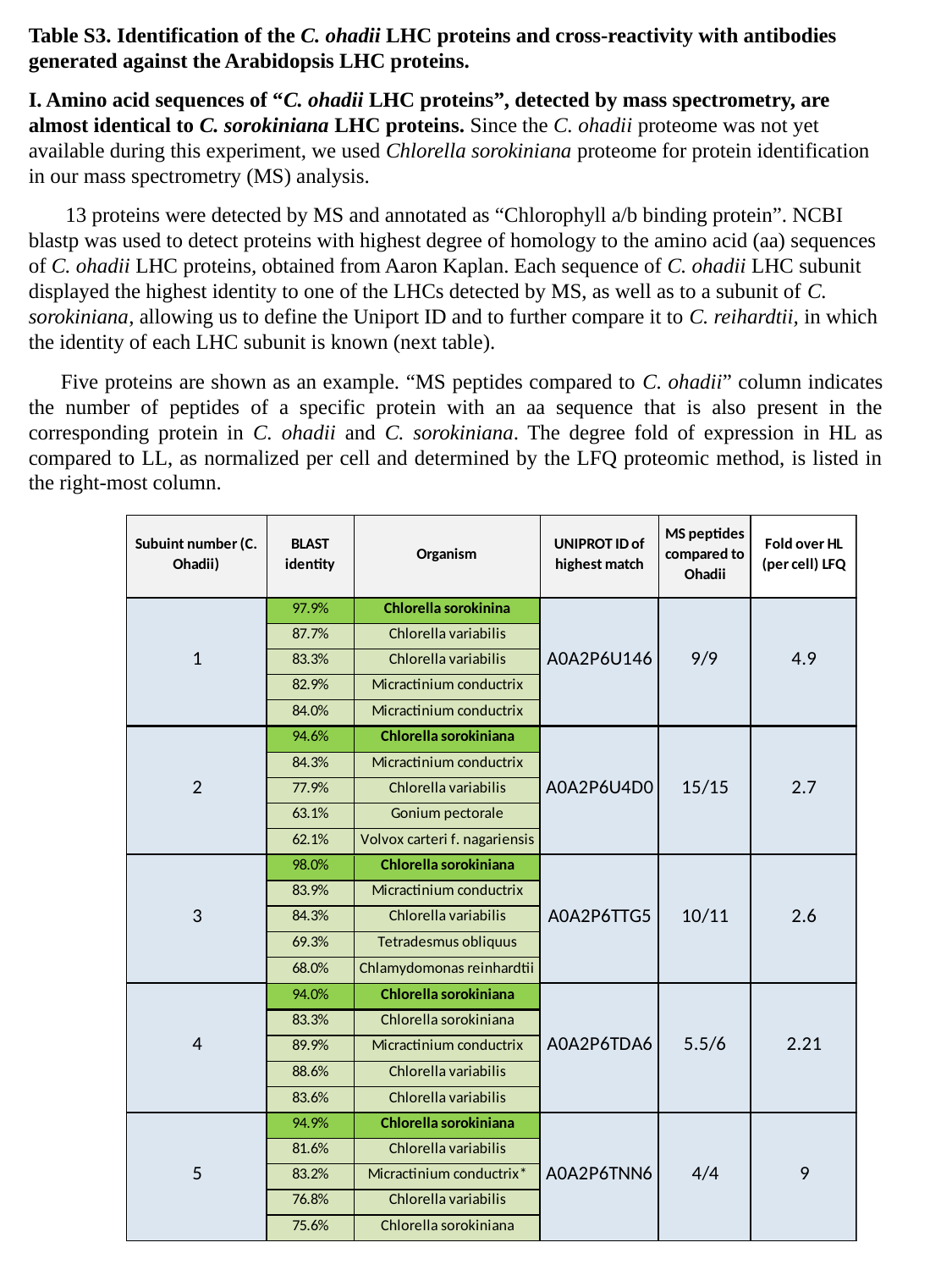

Table S3. Identification of the C. ohadii LHC proteins and cross-reactivity with antibodies generated against the Arabidopsis LHC proteins.
I. Amino acid sequences of “C. ohadii LHC proteins”, detected by mass spectrometry, are almost identical to C. sorokiniana LHC proteins. Since the C. ohadii proteome was not yet available during this experiment, we used Chlorella sorokiniana proteome for protein identification in our mass spectrometry (MS) analysis.
 13 proteins were detected by MS and annotated as “Chlorophyll a/b binding protein”. NCBI blastp was used to detect proteins with highest degree of homology to the amino acid (aa) sequences of C. ohadii LHC proteins, obtained from Aaron Kaplan. Each sequence of C. ohadii LHC subunit displayed the highest identity to one of the LHCs detected by MS, as well as to a subunit of C. sorokiniana, allowing us to define the Uniport ID and to further compare it to C. reihardtii, in which the identity of each LHC subunit is known (next table).
 Five proteins are shown as an example. “MS peptides compared to C. ohadii” column indicates the number of peptides of a specific protein with an aa sequence that is also present in the corresponding protein in C. ohadii and C. sorokiniana. The degree fold of expression in HL as compared to LL, as normalized per cell and determined by the LFQ proteomic method, is listed in the right-most column.

### Slide 2
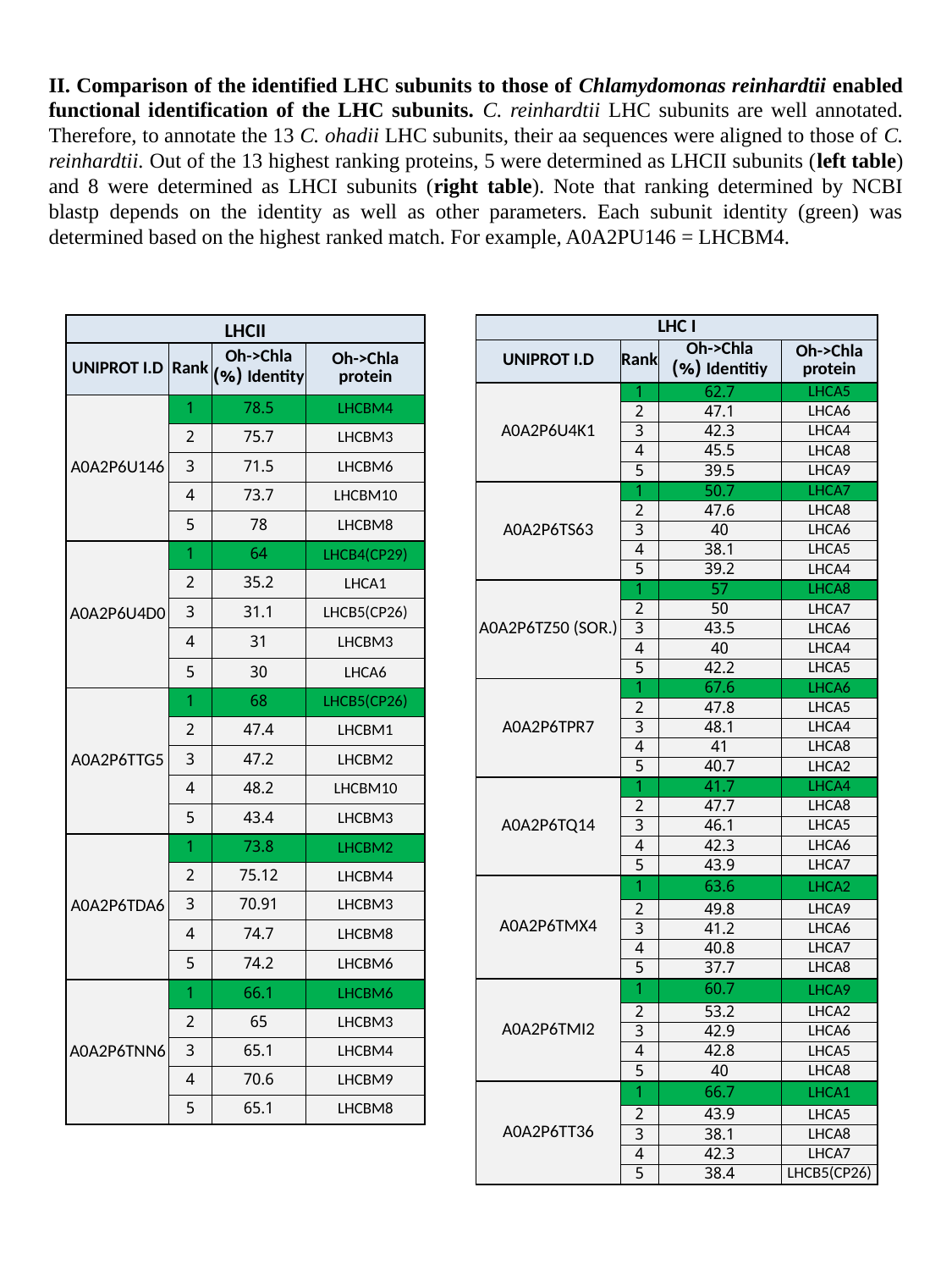

II. Comparison of the identified LHC subunits to those of Chlamydomonas reinhardtii enabled functional identification of the LHC subunits. C. reinhardtii LHC subunits are well annotated. Therefore, to annotate the 13 C. ohadii LHC subunits, their aa sequences were aligned to those of C. reinhardtii. Out of the 13 highest ranking proteins, 5 were determined as LHCII subunits (left table) and 8 were determined as LHCI subunits (right table). Note that ranking determined by NCBI blastp depends on the identity as well as other parameters. Each subunit identity (green) was determined based on the highest ranked match. For example, A0A2PU146 = LHCBM4.
| LHCII | | | |
| --- | --- | --- | --- |
| UNIPROT I.D | Rank | Oh->Chla Identity (%) | Oh->Chla protein |
| A0A2P6U146 | 1 | 78.5 | LHCBM4 |
| | 2 | 75.7 | LHCBM3 |
| | 3 | 71.5 | LHCBM6 |
| | 4 | 73.7 | LHCBM10 |
| | 5 | 78 | LHCBM8 |
| A0A2P6U4D0 | 1 | 64 | LHCB4(CP29) |
| | 2 | 35.2 | LHCA1 |
| | 3 | 31.1 | LHCB5(CP26) |
| | 4 | 31 | LHCBM3 |
| | 5 | 30 | LHCA6 |
| A0A2P6TTG5 | 1 | 68 | LHCB5(CP26) |
| | 2 | 47.4 | LHCBM1 |
| | 3 | 47.2 | LHCBM2 |
| | 4 | 48.2 | LHCBM10 |
| | 5 | 43.4 | LHCBM3 |
| A0A2P6TDA6 | 1 | 73.8 | LHCBM2 |
| | 2 | 75.12 | LHCBM4 |
| | 3 | 70.91 | LHCBM3 |
| | 4 | 74.7 | LHCBM8 |
| | 5 | 74.2 | LHCBM6 |
| A0A2P6TNN6 | 1 | 66.1 | LHCBM6 |
| | 2 | 65 | LHCBM3 |
| | 3 | 65.1 | LHCBM4 |
| | 4 | 70.6 | LHCBM9 |
| | 5 | 65.1 | LHCBM8 |
| LHC I | | | |
| --- | --- | --- | --- |
| UNIPROT I.D | Rank | Oh->Chla Identitiy (%) | Oh->Chla protein |
| A0A2P6U4K1 | 1 | 62.7 | LHCA5 |
| | 2 | 47.1 | LHCA6 |
| | 3 | 42.3 | LHCA4 |
| | 4 | 45.5 | LHCA8 |
| | 5 | 39.5 | LHCA9 |
| A0A2P6TS63 | 1 | 50.7 | LHCA7 |
| | 2 | 47.6 | LHCA8 |
| | 3 | 40 | LHCA6 |
| | 4 | 38.1 | LHCA5 |
| | 5 | 39.2 | LHCA4 |
| A0A2P6TZ50 (SOR.) | 1 | 57 | LHCA8 |
| | 2 | 50 | LHCA7 |
| | 3 | 43.5 | LHCA6 |
| | 4 | 40 | LHCA4 |
| | 5 | 42.2 | LHCA5 |
| A0A2P6TPR7 | 1 | 67.6 | LHCA6 |
| | 2 | 47.8 | LHCA5 |
| | 3 | 48.1 | LHCA4 |
| | 4 | 41 | LHCA8 |
| | 5 | 40.7 | LHCA2 |
| A0A2P6TQ14 | 1 | 41.7 | LHCA4 |
| | 2 | 47.7 | LHCA8 |
| | 3 | 46.1 | LHCA5 |
| | 4 | 42.3 | LHCA6 |
| | 5 | 43.9 | LHCA7 |
| A0A2P6TMX4 | 1 | 63.6 | LHCA2 |
| | 2 | 49.8 | LHCA9 |
| | 3 | 41.2 | LHCA6 |
| | 4 | 40.8 | LHCA7 |
| | 5 | 37.7 | LHCA8 |
| A0A2P6TMI2 | 1 | 60.7 | LHCA9 |
| | 2 | 53.2 | LHCA2 |
| | 3 | 42.9 | LHCA6 |
| | 4 | 42.8 | LHCA5 |
| | 5 | 40 | LHCA8 |
| A0A2P6TT36 | 1 | 66.7 | LHCA1 |
| | 2 | 43.9 | LHCA5 |
| | 3 | 38.1 | LHCA8 |
| | 4 | 42.3 | LHCA7 |
| | 5 | 38.4 | LHCB5(CP26) |

### Slide 3
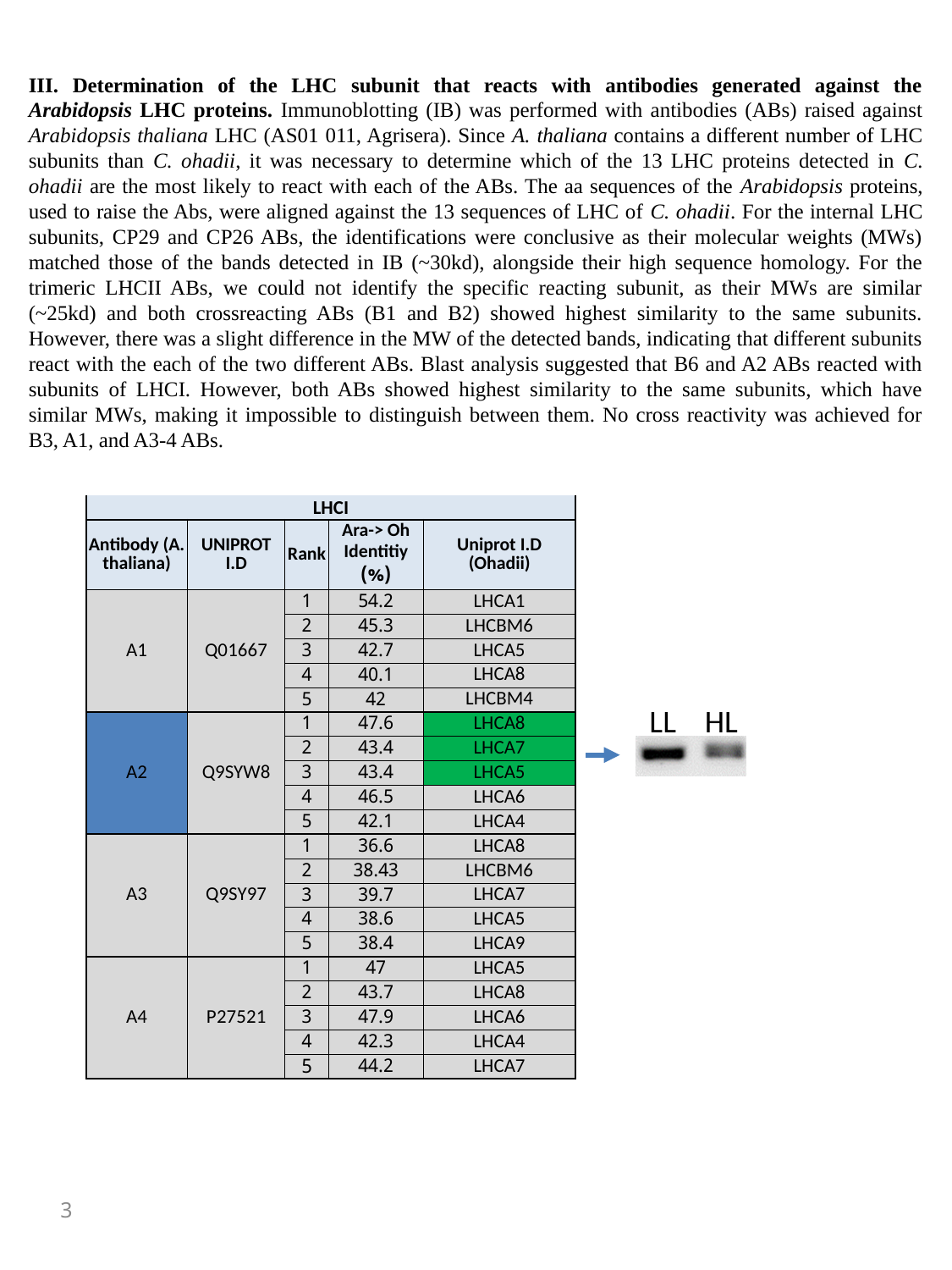

III. Determination of the LHC subunit that reacts with antibodies generated against the Arabidopsis LHC proteins. Immunoblotting (IB) was performed with antibodies (ABs) raised against Arabidopsis thaliana LHC (AS01 011, Agrisera). Since A. thaliana contains a different number of LHC subunits than C. ohadii, it was necessary to determine which of the 13 LHC proteins detected in C. ohadii are the most likely to react with each of the ABs. The aa sequences of the Arabidopsis proteins, used to raise the Abs, were aligned against the 13 sequences of LHC of C. ohadii. For the internal LHC subunits, CP29 and CP26 ABs, the identifications were conclusive as their molecular weights (MWs) matched those of the bands detected in IB (~30kd), alongside their high sequence homology. For the trimeric LHCII ABs, we could not identify the specific reacting subunit, as their MWs are similar (~25kd) and both crossreacting ABs (B1 and B2) showed highest similarity to the same subunits. However, there was a slight difference in the MW of the detected bands, indicating that different subunits react with the each of the two different ABs. Blast analysis suggested that B6 and A2 ABs reacted with subunits of LHCI. However, both ABs showed highest similarity to the same subunits, which have similar MWs, making it impossible to distinguish between them. No cross reactivity was achieved for B3, A1, and A3-4 ABs.
| LHCI | | | | |
| --- | --- | --- | --- | --- |
| Antibody (A. thaliana) | UNIPROT I.D | Rank | Ara-> Oh Identitiy (%) | Uniprot I.D (Ohadii) |
| A1 | Q01667 | 1 | 54.2 | LHCA1 |
| | | 2 | 45.3 | LHCBM6 |
| | | 3 | 42.7 | LHCA5 |
| | | 4 | 40.1 | LHCA8 |
| | | 5 | 42 | LHCBM4 |
| A2 | Q9SYW8 | 1 | 47.6 | LHCA8 |
| | | 2 | 43.4 | LHCA7 |
| | | 3 | 43.4 | LHCA5 |
| | | 4 | 46.5 | LHCA6 |
| | | 5 | 42.1 | LHCA4 |
| A3 | Q9SY97 | 1 | 36.6 | LHCA8 |
| | | 2 | 38.43 | LHCBM6 |
| | | 3 | 39.7 | LHCA7 |
| | | 4 | 38.6 | LHCA5 |
| | | 5 | 38.4 | LHCA9 |
| A4 | P27521 | 1 | 47 | LHCA5 |
| | | 2 | 43.7 | LHCA8 |
| | | 3 | 47.9 | LHCA6 |
| | | 4 | 42.3 | LHCA4 |
| | | 5 | 44.2 | LHCA7 |
LL HL
3

### Slide 4
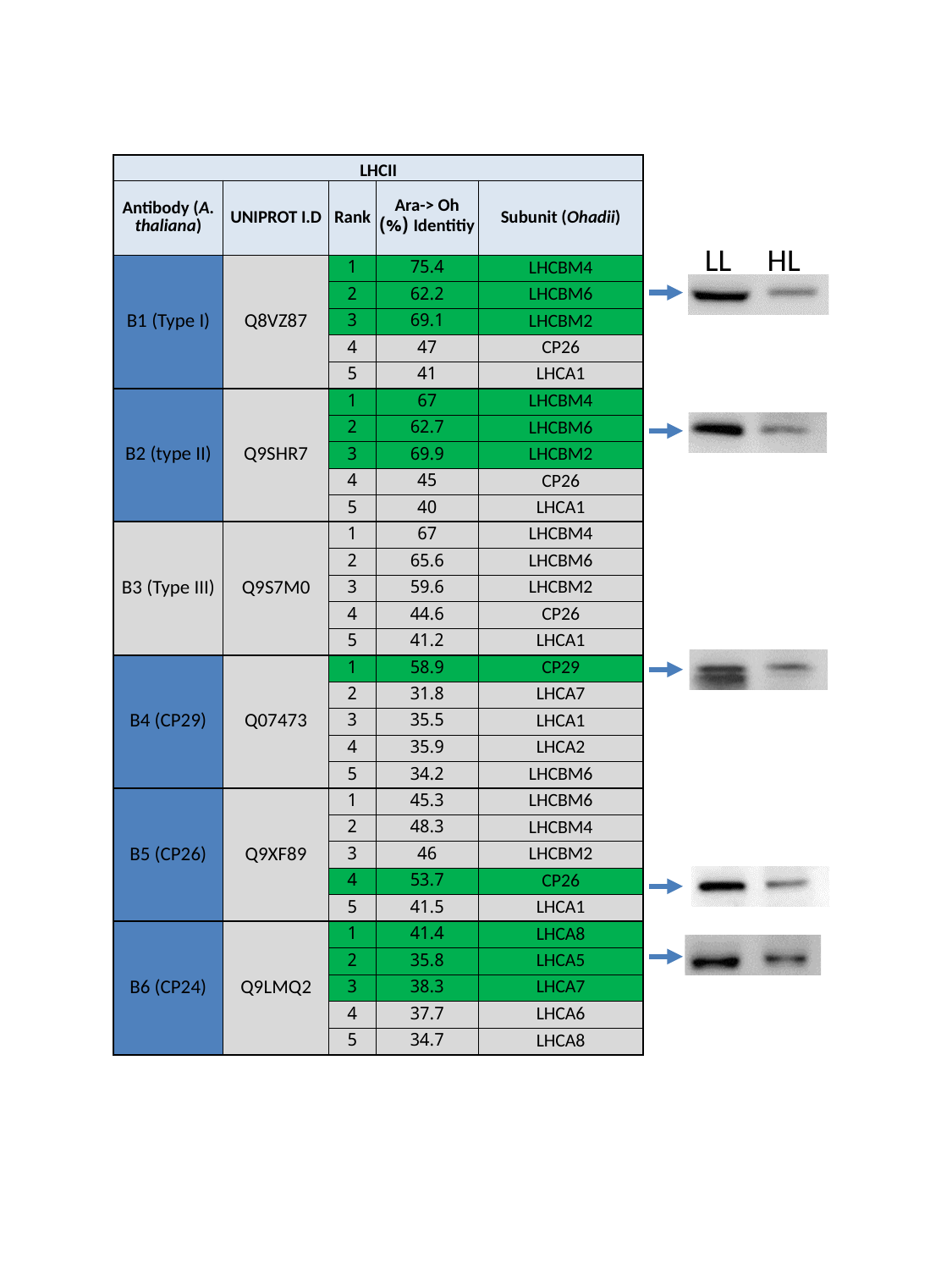

| LHCII | | | | |
| --- | --- | --- | --- | --- |
| Antibody (A. thaliana) | UNIPROT I.D | Rank | Ara-> Oh Identitiy (%) | Subunit (Ohadii) |
| B1 (Type I) | Q8VZ87 | 1 | 75.4 | LHCBM4 |
| | | 2 | 62.2 | LHCBM6 |
| | | 3 | 69.1 | LHCBM2 |
| | | 4 | 47 | CP26 |
| | | 5 | 41 | LHCA1 |
| B2 (type II) | Q9SHR7 | 1 | 67 | LHCBM4 |
| | | 2 | 62.7 | LHCBM6 |
| | | 3 | 69.9 | LHCBM2 |
| | | 4 | 45 | CP26 |
| | | 5 | 40 | LHCA1 |
| B3 (Type III) | Q9S7M0 | 1 | 67 | LHCBM4 |
| | | 2 | 65.6 | LHCBM6 |
| | | 3 | 59.6 | LHCBM2 |
| | | 4 | 44.6 | CP26 |
| | | 5 | 41.2 | LHCA1 |
| B4 (CP29) | Q07473 | 1 | 58.9 | CP29 |
| | | 2 | 31.8 | LHCA7 |
| | | 3 | 35.5 | LHCA1 |
| | | 4 | 35.9 | LHCA2 |
| | | 5 | 34.2 | LHCBM6 |
| B5 (CP26) | Q9XF89 | 1 | 45.3 | LHCBM6 |
| | | 2 | 48.3 | LHCBM4 |
| | | 3 | 46 | LHCBM2 |
| | | 4 | 53.7 | CP26 |
| | | 5 | 41.5 | LHCA1 |
| B6 (CP24) | Q9LMQ2 | 1 | 41.4 | LHCA8 |
| | | 2 | 35.8 | LHCA5 |
| | | 3 | 38.3 | LHCA7 |
| | | 4 | 37.7 | LHCA6 |
| | | 5 | 34.7 | LHCA8 |
LL HL
